# Identification of two β-cell subtypes by 7 independent criteria

**DOI:** 10.1101/2023.01.31.526222

**Authors:** Erez Dror, Luca Fagnocchi, Vanessa Wegert, Stefanos Apostle, Brooke Grimaldi, Tim Gruber, Ilaria Panzeri, Steffen Heyne, Kira Daniela Höffler, Victor Kreiner, Reagan Ching, Tess Tsai-Hsiu Lu, Ayush Semwal, Ben Johnson, Parijat Senapati, Adelheid M. Lempradl, Dustin Schones, Axel Imhof, Hui Shen, John Andrew Pospisilik

## Abstract

Despite the recent explosion in surveys of cell-type heterogeneity, the mechanisms that specify and stabilize highly related cell subtypes remain poorly understood. Here, focusing initially on exploring quantitative histone mark heterogeneity, we identify two major sub-types of pancreatic β-cells (β_HI_ and β_LO_). β_HI_ and β_LO_ cells differ in their size, morphology, cytosolic and nuclear ultrastructure, transcriptional output, epigenomes, cell surface marker, and function. Importantly, β_HI_ and β_LO_ cells can be FACS separated live into CD24^+^ (β_HI_) and CD24^-^ (β_LO_) fractions. From an epigenetic viewpoint, β_HI_-cells exhibit ∼4-fold higher levels of H3K27me3, more compacted chromatin, and distinct chromatin organization that associates with a specific pattern of transcriptional output. Functionally, β_HI_ cells have increased mitochondrial mass, activity, and insulin secretion both *in vivo* and *ex vivo*. Critically, *Eed* and *Jmjd3* loss-of-function studies demonstrate that H3K27me3 dosage is a significant regulator of β_HI_ / β_LO_ cell ratio *in vivo,* yielding some of the first-ever specific models of β-cell sub-type distortion. β_HI_ and β_LO_ sub-types are conserved in humans with β_HI_-cells enriched in human Type-2 diabetes. These data identify two novel and fundamentally distinct β-cell subtypes and identify epigenetic dosage as a novel regulator of β-cell subtype specification and heterogeneity.

**Highlights:**

- Quantitative H3K27me3 heterogeneity reveals 2 common β-cell subtypes
- β_HI_ and β_LO_ cells are stably distinct by 7 independent sets of parameters
- H3K27me3 dosage controls β_HI_ / β_LO_ ratio in vivo
- β_HI_ and β_LO_ cells are conserved in humans and enriched in Type-2 diabetes

## Introduction

β-cells are the sole providers of insulin in the body, acting to optimize nutrient uptake and storage, and to prevent hyperglycemia. During development, β-cells differentiate through progressive activation of transcription factor-directed gene networks and undergo functional maturation during early post-natal life (Salinno et al., 2019; Stolovich-Rain et al., 2015). Adult β-cells are highly specialized, quiescent and represent one of the longest-lived cell types in the body, averaging ∼30-40 years in elderly humans (Arrojo et al., 2019; Cnop et al., 2011). β-cells therefore rely on specific epigenetic systems to stabilize and maintain cell identity over expansive time scales (Dhawan et al., 2011; Lu et al., 2018). A relative loss of functional pancreatic β-cell mass results in diabetes.

Significant cell-to-cell heterogeneity has been observed within the β-cell compartment since at least 1960 (Hellerstrom et al., 1960). Early studies found heterogeneity in glucose thresholds, calcium handling, and insulin secretion (Kiekens et al., 1992; Salomon and Meda, 1986), observations that have been confirmed using advanced optical and genetic tools (Johnston et al., 2016; Salem et al., 2019). Recent applications of specialized molecular tools and mouse models allowed the identification of factors that affect β-cell heterogeneity including the maturation marker Cfap126 that identified a primary maturation gradient (Bader et al., 2016), virgin cells (van der Meulen et al., 2017) and immune evading subsets (Rui et al., 2017). The recent wide-spread adoption of single-cell technologies has re-focused attention on the origins, architecture and potential therapeutical relevance of β-cell heterogeneity, and a range of sub-states or sub-types have been proposed (Chiou et al., 2021; Dorrell et al., 2016; Salinno et al., 2021; Segerstolpe et al., 2016; van der Meulen et al., 2017; Xin et al., 2018). Despite the intense research, however, the field has yet to assemble a universal framework for understanding β-cell sub-types and sub-states (Mawla and Huising, 2019; Wang and Kaestner, 2019).

Challenges to establishing such a framework include a relative over-reliance on single-cell genomic techniques. These technologies overall represent shallow snapshots of the transcriptome, are almost entirely biased towards the active epigenome, involve numerous bioinformatic assumptions, and fail to distinguish ‘cell-state’ from ‘cell-type’ heterogeneity. Cell-state, which is primarily characterized by transient, periodic or progressive temporal dynamics, comprise a substantial fraction of observed heterogeneity. Cell-state heterogeneity described in β-cells includes for instance circadian oscillations (Szabat et al., 2009), transcriptional bursting (e.g. at *Ins2* (Farack et al., 2019)), transcriptional noise (Enge et al., 2017), cell cycle (Dagogo-Jack and Shaw, 2018), maturation (Qiu et al., 2017), stress (Cigliola et al., 2016; Xin et al., 2018) and aging (Enge et al., 2017). These dimensions, which are effectively studied by imaging, are to definitively parse and regress out of single cell genomics data. Finally, use of different transgenic reporter systems across studies has added an additional challenge when examining and comparing all heterogeneity-focused data. *Cre* recombinase for instance is known to trigger ER stress and thus generate artificial heterogeneity signals (Rosenbaum et al., 2014; Xiao et al., 2012); similarly, inherently imperfect reporter expression leaves uncertainty as to whether heterogeneity is being accurately represented and/or artificially generated (Estall and Screaton, 2020).

Here, using a range of reporter-independent approaches, we find that the primary axis of β-cell heterogeneity is epigenetically-defined, and that it separates β-cells into two fundamentally distinct cell types (β_HI_ and β_LO_) with distinct morphology, cytosolic and nuclear ultrastructure, transcriptome output, epigenome configuration, and function. In healthy adult mice, β_HI_ and β_LO_ cells comprise >90 of β-cells. They are present at an approximate 1:4 ratio (β_HI_/β_LO_) from lactation through to old age and can be FACS sorted live into CD24+ and CD24− populations. β_HI_ and β_LO_ cells both exhibit robust proliferation *in vivo* and *in vitro*. β_HI_ cells appear to proliferate faster at baseline and their relative number are increased upon chronic high-fat diet (HFD). H3K27me3 dosage controls β_HI_/β_LO_ ratio *in vivo*, with conditional heterozygosity of the polycomb repressive complex 2 (PRC2) core subunit Eed and the histone de-methylase Jmjd3, generating equal and opposite cell ratio skewing. Eqaully important, we demonstrate that β_HI_ and β_LO_ cells are conserved in humans and that β_HI_ cells are enriched in Type 2 diabetes. These data identify two major β-cell sub-types, and identify epigenetic dosage as a novel and potentially targetable mechanism controlling β-cell compartment heterogeneity.

## Results

### Two common and epigenetically distinct β-cell sub-types

Historically, cell types were distinguished based on histopathological and nuclear differences, features that reflect stable differences in epigenome configuration (Rehimi et al., 2016; Skinner and Johnson, 2017; Stephens et al., 2019). To directly measure such epigenetic heterogeneity in β-cells, we quantified total histone modification levels at single-cell resolution using FACS, including total H3K4me3 (active promoters), H3K27ac (active cis-regulatory elements), H3K36me3 (transcribed gene-bodies), H3K9me3 (silent constitutive heterochromatin), and H3K27me3 (polycomb-associated heterochromatin). To avoid confounding potential artefacts associated with transgenic reporters, we performed antibody-based purification of freshly isolated, dissociated and fixed islet cells isolated from wildtype mice. We used insulin as a positive selection marker for all β-cells, and gated out cells that stained for CD45 (immune), CD31 (endothelial), SST (delta), GCG (alpha), and PP. Most histone marks showed robust and uniform immunoreactivity across all β-cells (Figure 1A; cell gating strategy, Figure S1A). Surprisingly, the signal for H3K27me3 appeared bimodal, suggesting two epigenetically distinct sub-populations (-LO vs -HI; Figures 1A, 1B). Averaged across independent biological replicates, -HI cells had a ∼4.5-fold higher H3K27me3 mean fluorescence intensity (MFI) than -LO cells (Figure S1B). Imaging-flow-cytometry validated that the H3K27me3 signal in both -HI and -LO cells was nuclear in origin and ruled out cell-doublets, poly-nucleated cells, and cytosolic immunoreactivity as potential confounding sources for the -HI signal (Figure 1C). H3K27me3-HI and -LO populations were consistently observed across experiments, animals, ages, and within islets of both males and females (Figure 1D; Figure S1C). In female β-cells, no difference in H3K27me3 immunoreactivity was observed on the silent X-chromosome (Barr body), further highlighting the specificity of the H3K27me3 differences (Figures S1C, D). Importantly, -HI and -LO cells were found in all islets of all sizes (Figures S1C, E) arguing against inter-islet differences as the source of observed epigenetic signature. The H3K27me3 signal was validated using two independent antibodies (Figure 1A, monoclonal; and Figure S1F, monoclonal vs. polyclonal) and against β-cell-specific Eed/PRC2 knockout (KO) mouse islets that are deficient in H3K27me3 (βEedKO; Figure S1G). Parallel analyses of pancreatic islet ɑ-, δ- and PP-cells suggested that the H3K27me3 signature was specific to β-cells (Figure S1H). Thus, β-cells exist in two common populations distinct in their H3K27me3 levels.

**Figure 1.**
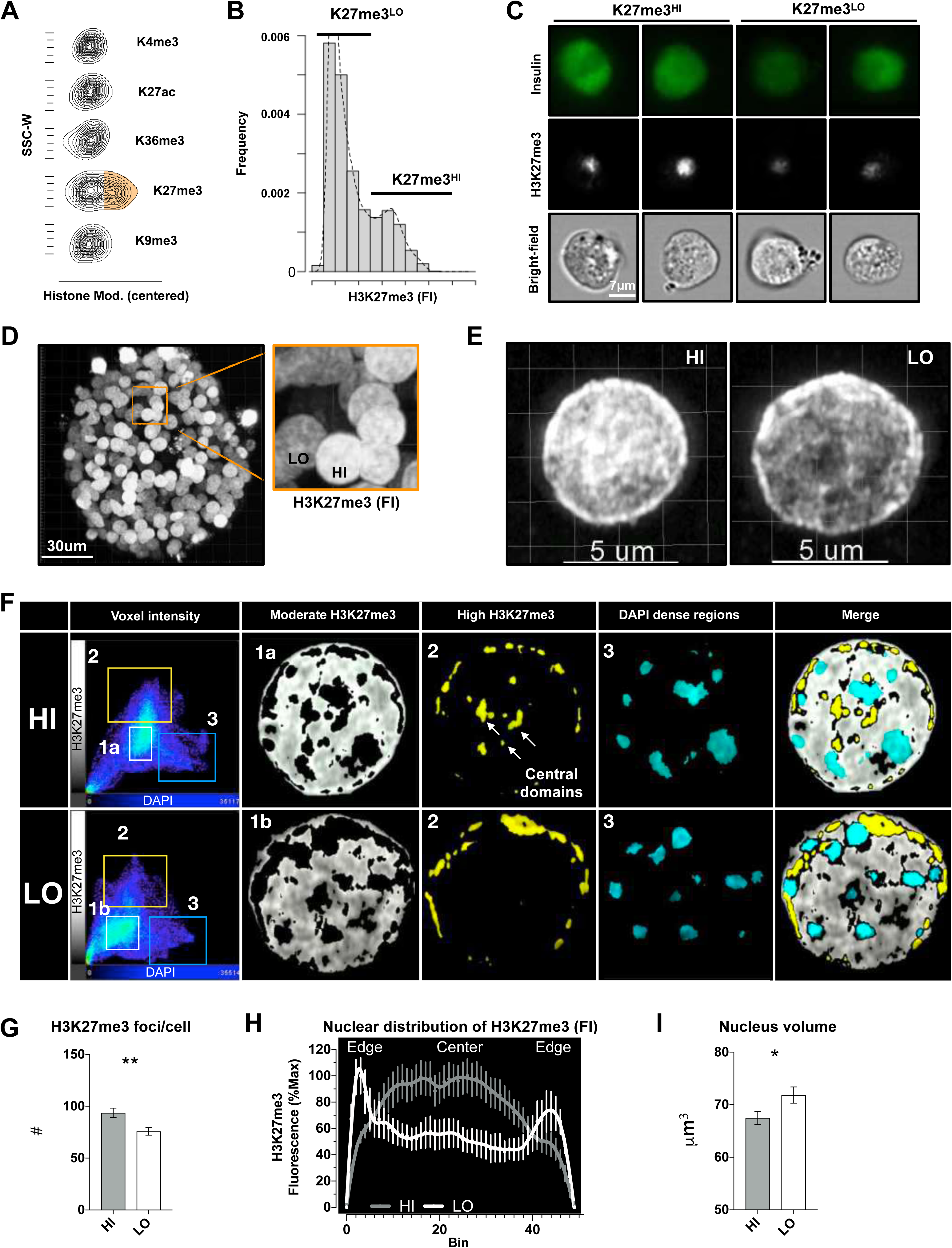
Two epigenetically distinct pancreatic β-cell sub-types. A. Representative contour plots of the centered intensities of the stated histone modifications in β-cells isolated from individual mice (image representative of n=6 mice from 3 experiments). B. Representative distribution plots of H3K27me3 staining fluorescent intensities (FI) in insulin positive β-cells (representative of 5 experiments, n=4 mice each). C. Representative ImageStream analysis of dispersed, fixed single- β-cells of islets isolated from individual mice. The different panels show the immunostaining against insulin (up) H3K27me3 (middle), and bright field image (bottom) of the same β-cells (representative of 2 experiments). D. Representative 3D reconstruction of one pancreatic islet isolated from male mice, immunostained against H3K27me3, and a zoomed in image of adjacent insulin positive H3K27me3 HI and LO nuclei (representative of 3 experiments). E. 3D reconstruction of high-resolution confocal imaging from H3K27me3-HI (left) or -LO (right) sorted β-cells; one representative image out of 60 nuclei from n=4 mice. F. Representative voxel intensities and co-localizations of H3K27me3 and DAPI in one z-plane of each of the nuclei imaged in E. Groupings of voxels was done according to their DAPI and H3K27me3 intensities (left panel). Group 1 represent low\moderate intensity voxels, are localized in the nuclear interior and are shifted when comparing -HI and -LO cells (1a and 1b). H3K27me3 high intensity voxels are in group 2 (yellow) and are localized in the nuclear periphery of both nuclei with addition of central domains in the H3K27me3-HI nucleus. DAPI high voxels are in group 3 that is unchanged. G. Bar plot representation of the mean of numbers of H3K27me3 foci per nucleus of HI/LO β-cells isolated from 4 individual mice. Assessed by automated quantification of high-resolution images of 67 (HI) and 63 (LO) single nuclei. **= unpaired t-test, *p*-value<0.01. Error bars are mean ± SEM. H. Line plot of the averaged H3K27me3 intensities across the center optical plane (binned) of HI/LO sorted β-cell nuclei. Signal is normalized per cell. I. Bar plot representation of the Mean of the nucleus volumes of HI/LO sorted β-cells as assessed after reconstructing DAPI positive z-stacks and measuring the DAPI positive volume (analysis of high-resolution imaging of the 67 or 63 nuclei of single cells isolated from 4 individual mice). *= unpaired t-test, *p*-value <0.05. Error bars are mean ± SEM.

Next, we used super-resolution confocal microscopy to validate the findings and test for differences in nuclear morphology (Figures 1E-I). Imaging and analysis revealed clear distinctions between the -HI and -LO cells, in that H3K27me3-HI cells contained more H3K27me3-foci (Figures 1F, G). H3K27me3-foci in other systems have been associated with compacted Polycomb-silenced genomic regions (Boettiger et al., 2016). Further, whereas -LO cells showed H3K27me3 staining primarily at the transcriptionally silent nuclear periphery, -HI cell H3K27me3 was enriched in the active nuclear interior (Geyer et al., 2011) (Figure 1F box 2, Figure 1H, and Figure S1I). Consistent with the role of H3K27me3 in chromatin silencing and compaction (Eskeland et al., 2010), nuclei were ∼5 µm^3^ smaller (on average) in the -HI relative to -LO cells (Figure 1I). Thus, pancreatic β-cells exist in two populations based on H3K27me3 level, chromatin organization, and nuclear compaction.

### H3K27me3-HI cells are transcriptionally distinct and express cell surface CD24

To determine if the H3K27me3 difference between -HI and -LO β-cells translated into stable differences in transcriptome output, we FACS sorted -HI and -LO β-cells (INS+ but GCG^-^/SST^-^/PP^-^/CD31^-^/CD45^-^) from eight individual wildtype mice across two age groups (4 or 10 weeks old; Figure 2A) and performed RNA-seq. By principal component analysis (PCA), -HI and -LO H3K27me3 status separated on PC1, indicating that stable and reproducible transcriptome differences exist between the -HI and -LO β-cells, and that these are maintained from weaning (4 weeks) into adulthood (Figure 2B). Genes differentially expressed between -HI and -LO cells were enriched for a set of near-silent or poised genes known as bivalent genes (Lu et al., 2018). Intriguingly, these H3K27me3- dependent genes were upregulated in -HI β-cells (Figure 2C) suggesting they may be preferentially transcribed in one of the two subpopulations. Notably, this set of differentially regulated genes included alternate islet endocrine lineages factors (*Ppy*, *Gcg*, and *Sst*), as well as heterogeneity and plasticity markers (*Arx*, *Etv1*, *Gpx3* and *Rbp4*). Importantly, two differentially expressed genes coded for cell surface proteins (*Slc23a4* and *Cd24a*). Slc23a4 is annotated as a pseudogene in humans. We obtained antibodies to CD24 (the protein product of *CD24a*) and performed anti-CD24 antibody titrations and FACS analysis to test if -HI and -LO β-cell populations could be distinguished based on cell surface staining. Importantly, co-staining of live cells isolated from β-cell reporter mouse with CD24 and H3K27me3 refined the partially overlapping -HI and -LO subsets into two separate β-cell populations (Figure 2D, gating strategy in Figure S2A), with ∼20% of all INS^+^ β-cells being CD24-positive (CD24^+^) and ∼80% CD24− negative (CD24**^-^**, Figure 2D). CD24^+^ cells showed higher levels of H3K27me3 (Figures 2D, E). Noteworthy, we also observed rare INS^+^ cells (∼1%) with extremely high CD24 levels (CD24^high^ in Figure S2B, left panel). These CD24^high^ cells were SST^+^ (Figure S2B right panel) and are consistent with prior studies showing strong δ-cell expression of CD24 (Berthault et al., 2020). These rare double-hormone positive (INS^+^/SST^+^) cells were excluded from all further analyses.

**Figure 2.**
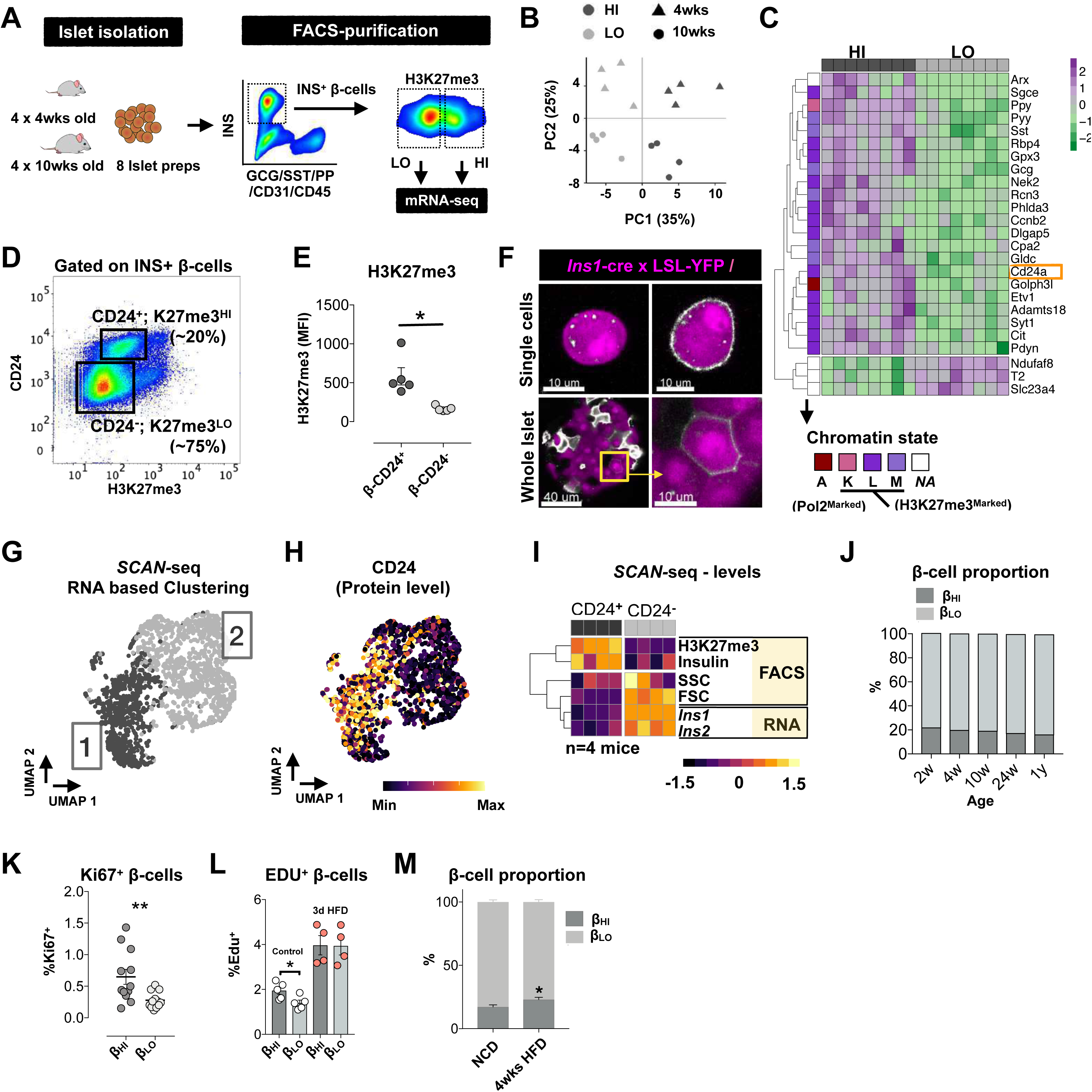
H3K27me3-HI cells are transcriptionally distinct and express cell surface CD24. A. Schematic of the experimental plan. Eight biological replicates of H3K27me3 HI/LO β-cells were isolated from four 4-week-old or four 10-week-old wildtype mice. One thousand H3K27me3 HI/LO cells were sorted from each mouse and low input RNA extraction and mRNA-seq was performed. B. PCA RNA-seq signals across the β-cells from young and adult mice used in the screening study. Each data point, shown as a triangle or a circle, represents the transcriptome of HI (dark gray) or LO (light gray) β-cells isolated from individual 4 (triangles) or 10 (circles) weeks old mice. total of n=8 mice. C. Heatmap of the differentially expressed genes between H3K27me3-HI/LO murine β-cells, and their chromatin-states as previously annotated (Lu et al., 2018). Log(normalized counts), z-scored per row. D. Representative example of CD24 expression versus H3K27me3 intensities in β-cells isolated from 10-week-old wildtype mice. representative of n=5 experiments. E. H3K27me3 mean fluorescence intensities (MFI) in CD24^-/+^ β-cells; each dot represents a population from an individual mouse. Paired t-test, * represent *p*-value<0.05. n=5 experiments. Error bars are mean± SEM. F. Representation of the heterogeneity in CD24 expression in live single β-cells or whole islets isolated from β-cell reporter mouse line (YFP is expressed upon Ins1-promoter driven CRE expression). G. UMAP visualization of sorted mouse β-cells that underwent *SCAN-seq* protocol. Colors represent the two major clusters of β-cells. n=2,156 cells H. UMAP map overlaid with the FACS-recorded levels of CD24 protein of each cell. I. Heatmap representation of SCAN-seq-scaled and averaged values (FACS-recorded intensities of the depicted parameters or RNA expression levels; Z-scored per row) from single β-cells negative or positive for CD24 from n=4 individual mice (columns). J. Representation of the proportion of H3K27me3-HI\CD24^+^ β cells through the life-span of mice. 8-12 mice per age group from n=4 experiments. Error bars are mean± SEM K. Representation of the proliferating cell fraction of H3K27me3-HI\CD24^+^ and H3K27me3-LO\CD24^-^ β cells. Paired t-test, * represent *p*-value<0.05. Each dot represents one mouse, 12 mice from a total n=4 experiments. Error bars are mean± SEM L. Representation of the proliferation in the H3K27me3-HI\CD24^+^ or the H3K27me3-LO\CD24^-^ β cell compartment during 3 days of normal chow diet (control) or high fat diet (HFD) feeding. Mice were injected with Edu once per day. Paired t-test, * represent *p*-value<0.05. Each dot represents one mouse, 4-5 independent mice. Error bars are mean± SEM M. Representation of the proportion of H3K27me3-HI\CD24^+^ β-cells upon 4 weeks of high fat diet feeding. Unpaired t-test, * represent *p*-value<0.05. 10-11 mice per treatment group from n=3 experiments. Error bars are mean± SEM.

We validated the CD24 surface stain in several ways. By using an *Ins1-*YFP reporter mouse (*Ins1*-cre x LSL-YFP) and confocal imaging, we found CD24 expression was restricted to a subset of live β-cells, and determined that CD24 protein expression is cell membrane specific in single cells (Figure 2F upper panel) and in whole islets (Figure 2F lower panel, note the dim labeling compared to the YFP negative, CD24^high^ delta cells). Importantly, live-sorted Ins1-YFP^+^/CD24^+^ double-positive cells also showed higher H3K27me3 (Figure S2C, left) and nuclear compaction (Figure S2C, right), indicating that CD24^+^ and H3K27me3-HI β-cells are largely the same. Specificity of the CD24 antibody was confirmed using β-cells from CD24 knockout mice (Figure S2D). Thus, CD24 surface expression discriminates H3K27me3-HI and H3K27me3-LO β-cells.

To associate these findings with transcriptional heterogeneity, we modified one of the most sensitive single-cell (sc) RNA-seq protocols available, CELseq2 (Ziegenhain et al., 2017), to enable concomitant quantification of cell *<u>S</u>*urface (CD24), *<u>C</u>*ytoplasmic (Insulin) *<u>a</u>*nd *<u>N</u>*uclear (H3K27me3) protein epitopes (all at single-cell resolution) and applied this to purified INS^+^ β-cells isolated from wildtype mice. The new ‘*SCAN*-seq’ method is outlined in detail in the methods. As reported elsewhere (Chiou et al., 2021; Xin et al., 2018), transcriptome-based UMAP projection across all measured β-cells identified two major clusters (Figure 2G), with additional clusters emerging as the threshold stringency for within-cluster heterogeneity was increased (Figure S2E). Consistent with the data above, one of the two main clusters exhibited elevated expression of H3K27me3 marked genes (Figure S2F). Visualization of quantitative protein measures (FACS-based) onto the transcriptome-based UMAP showed that cells with CD24 surface-staining and high H3K27me3 levels clustered together to form one of the two major clusters (Cluster 1, Figures 2G -I). Also consistent with the data above, cells of the CD24^+^ transcriptomic cluster were smaller (lower FSC) and showed distinct granularity (SSC) relative to the CD24^-^ cluster. Notably, these observations would likely be overlooked by conventional scRNAseq due to the low level of *Cd24a* mRNA expression. The CD24^+^ transcriptomic cluster showed lower *Ins1* and *Ins2* mRNA counts despite higher insulin protein staining, highlighting the added value of *SCAN*-seq over conventional scRNA-seq. Thus, H3K27me3-HI and H3K27me3-LO β-cells can be separated by single-cell transcriptomics and CD24 expression.

### β_HI_ and β_LO_ cells

The preceding data demonstrate that CD24^+^/H3K27me3-HI and CD24^-^/H3K27me3-LO β-cells are distinct by bulk and single-cell transcriptomics, nuclear ultrastructure, nuclear and cell size, and total H3K27me3. The sub-types can be FACS-sorted live by their modest difference CD24 cell surface expression. They also demonstrate that H3K27me3-marked genes stratify a primary axis of transcriptional heterogeneity in the β-cell compartment (Figure 2C, Figure S2F). Importantly, reanalysis of published scRNA-seq datasets (Avrahami et al., 2020; Balboa et al., 2022; Pineros et al., 2020; Sachs et al., 2020; Xin et al., 2018) validated that the CD24/H3K27me3 axis is represented in the primary UMAP ‘dimensions’ of β-cell transcriptional heterogeneity reported across independent labs and studies (Figure S2F), and, that expression of H3K27me3-controlled genes separates β-cells into two primary clusters in both mice (Figure S2G) and humans (Figure S2H). We also validated the axis in human β-cell single nucleus (sn) ATACseq datasets (Chiou et al., 2021) which use chromatin accessibility as a measure of transcriptional potential (Figure S2I). Cell-state heterogeneity and in particular gradients of expression of the previously reported heterogeneity markers (where detectable), appear predominantly as gradients *within* CD24/H3K27me3 discordant clusters (UMAP matrices in Figure S2K, L). Thus, the CD24/H3K27me3 axis is evident in the primary dimension of β-cell heterogeneity across publicly available datasets.

Given the consistency across public datasets, and their separation based on morphological, nuclear, epigenetic transcriptional and cell-surface levels, we named these cells **β_HI_** (higher CD24; high H3K27me3; high chromatin compaction) and **β_LO_** cells. In our hands, β_HI_ and β_LO_ cells comprise ∼90-95% of all insulin protein positive pancreatic β-cells in adult mice. They are detectable from pre-weaning up to one year of age in mice (Figure 2J). Potentially important, we observe a progressive increase in H3K27me3 in both populations with age in mice (Figure S2J) as well as a very slow decline in β_HI_ / β_LO_ ratio. Based on Ki67 staining of freshly isolated islets, β_HI_ and β_LO_ cells both harbor proliferative capacity with a mild but significant increase in β_HI_ cells (Figure 2K). Importantly, proliferative capacity was validated *in vivo* using Edu-incorporation, data that also reveal that proliferative responsiveness in both cell sub-types upon 3-days of high fat diet (Figure 2L). And there was increase in β_HI_/β_LO_ cell type proportions upon chronic high fat feeding (4-weeks, Figure 2M). The majority of the murine β-cell compartment therefore comprises β_HI_ and β_LO_ cells.

### β_HI_ and β_LO_ cells exhibit distinct transcriptomes

Next, we separated β_HI_ and β_LO_ cells by FACS (CD24 and H3K27me3), increased replicate numbers, and performed bulk RNA-seq to enable differential expression analysis of entire transcriptome (active and silent). Despite strong transcriptional similarity (Figure S3A), we identified >2500 differentially expressed genes (Figures 3A-C). including those coding for mitochondrial and amide metabolic processes, oxidative phosphorylation, nuclear RNA processing factors, and interestingly, histone modification (Figure 3D). Consistent with their H3K27me3-HI phenotype, β_HI_ cells showed upregulation of *Ezh2,* the main H3K27me3-depositing methyltransferase. Relative to β_LO_ cells, β_HI_ cells also exhibited increased expression of the H3K27me3 demethylase *Kdm6b* (*Jmjd3*); the active mark ‘erasers’ *Hdac4/5*, the chromatin silencers *Cbx4*, *Suv420h2*, *Uhrf2*, and *Ehmt1/2*; and the 3D looping factors *Ctcf* and *Kmt2c*/*Mll3* (Figure 3E; blue, and Figure S3B). These data suggest that a persistent and complex network of chromatin regulation may exist to reinforce β_HI_ and β_LO_ cell differences. Importantly, we found no evidence for differences in the hallmark differentiation factors *Pdx1*, *Neurod1*, *Pax6, Nkx6.1*, *Mafa, Nkx2.2, Cfap126,* or *Cd81* (Figure 3E; beige), data that validated at the single cell level (Figure S2K, L). Modest opposing regulation however of the maturation factors *Ucn3* and *Rfx6* was observed, with *Rfx6* modestly up-regulated in β_HI_ cells (Figure 3E; red). Consistent with these findings, the Rfx6 binding motif was one of several motifs enriched at promoters of β_HI_ upregulated genes (Figures 3F and S3C). Also, in keeping with the *SCAN-*seq data (Figures 2G-I), β_HI_ cells showed modest decreases in *Ins1* and *Ins2* transcript levels (Ins1/2; Figure 3E, black, Figure S3D) despite clearly increased levels of insulin protein during FACS purification (Figure 3G). Thus, β_HI_ and β_LO_ are highly differentiated β-cells with distinct patterns of metabolic and chromatin regulatory gene expression.

**Figure 3.**
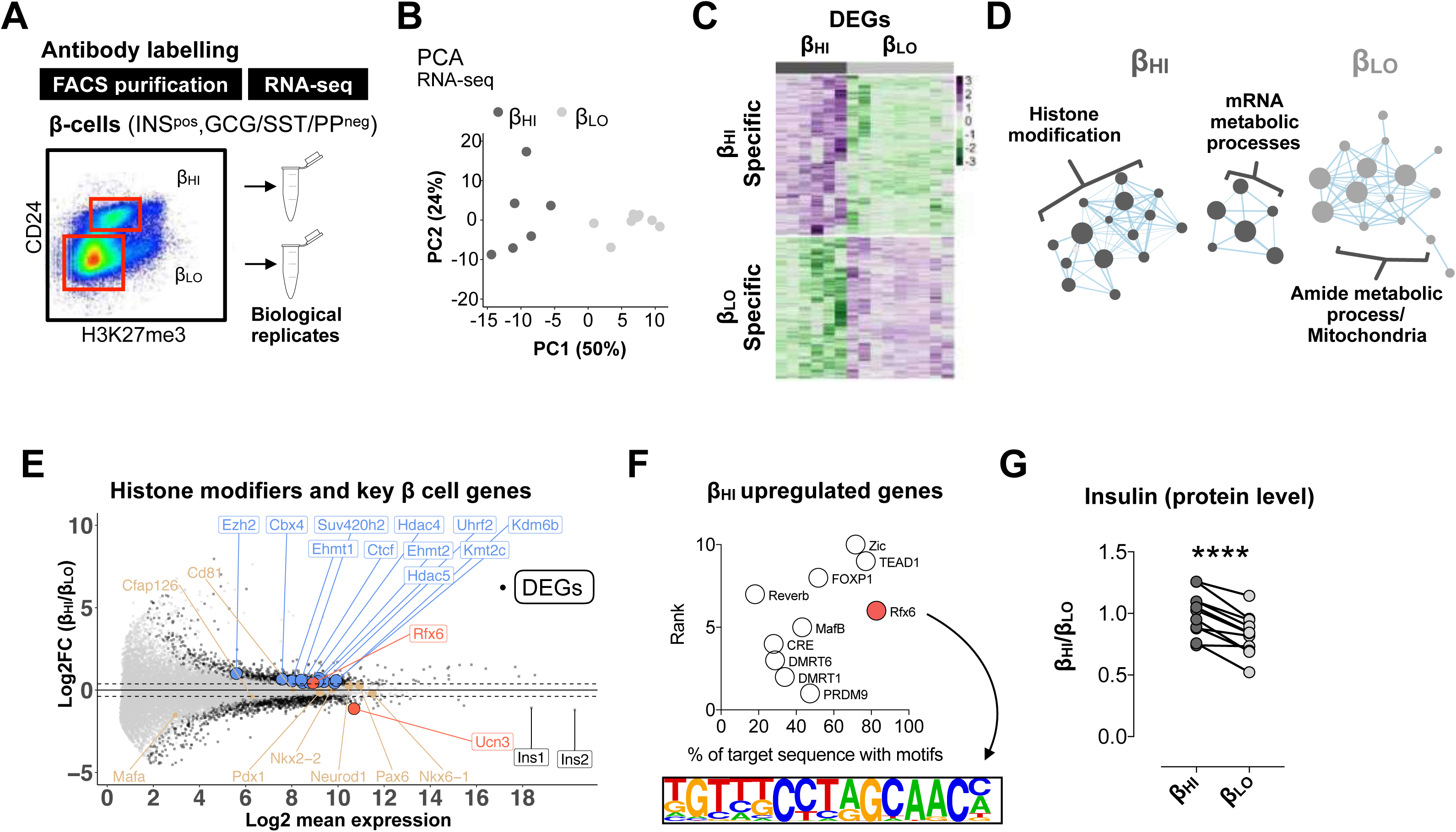
β_HI_ vs β_LO_ cells are functionally distinct and specialized. A. Schematic of the experimental plan. Two dimensions, CD24 and H3K27me3 allows clean separation of β_HI_/ β_LO_ for RNA sequencing analysis. B. PCA of H3K27me3 RNA-seq signals, showing reproducible separation of β_HI_ and β_LO_ β-cells, each dot represents one biological replicate from 3 independent experiments. C. Clustered heatmap representation of the log(normalized) expression of all differentially expressed genes (n=∼2500) across all replicates, Z-score was calculated per gene (row). D. A Cytoscape plot of GSEA pathways represents the β_HI_ (dark gray) or β_LO_ (light gary) enriched gene sets. Dot size is proportional to the false discovery rate *q*-value. E. MA plot showing the fold change in expression generated by comparing β_HI_ over β_LO_ β-cells. Black dots represent significantly deregulated genes, that are also boxed when labeled and highlighted (histone modifiers-blue; genes associated with β-cells and their maturation-red/beige; Ins1/2 genes -black). Black or Boxed genes are statistically significant (*P*-value adjusted for multiple testing < 0.05, with fold change cutoff of 1.33). F. Top 10 significant transcription factor motifs enriched within +/− 2kb from TSS of upregulated genes in β_HI_ cells. Rfx6 transcription factor and its binding motif are highlighted. G. Fold increase in insulin protein levels of β_HI_ cells. Connected dots represent cells from each of the types isolated from an individual mouse. **** = paired t-test, *p*-value <0.0001.

### β_HI_ and β_LO_ cells exhibit distinct epigenomes

To better understand the observed differences in nuclear ultrastructure and H3K27me3 levels, we FACS purified β_HI_ and β_LO_ cells from wildtype mice and performed H3K27me3 ChIP-seq on three paired independent biological replicates. As with their transcriptomes, β_HI_ and β_LO_ cell H3K27me3 profiles were strongly correlated (Figure 4A and Figure S4A). In keeping with published literature (Boyer et al., 2006; Lu et al., 2018; Margueron and Reinberg, 2011), we observed H3K27me3 enriched at focal regions across the genome, and across broad transcriptionally silent domains containing developmental genes, such as the Hox clusters and imprinted loci (Figures 4A, B and Figures S4B, C). PCA separated β_HI_ and β_LO_ cells on the first principal component, indicating both high quality data and reproducible differences (Figure 4C). H3K27me3 levels were unchanged at broad domains (Figure 4B and Figures S4B, C). Rather, differential H3K27me3 deposition was enriched at genic promoters and transcriptional start sites (TSS; Figure 4D and Figure S4D), suggesting H3K27me3 might underpin global cis-regulatory differences in gene expression between the two cell types. To test this idea, we called differential H3K27me3 enrichment at TSSs (Figure 4E, Figure S4E) and explored the relationship with transcriptome output. We detected ∼5200 differential peaks across ∼4750 annotated TSSs, and ∼1550 unique genes. Approximately 80% of these genes showed relative H3K27me3 enrichment in β_LO_ β-cells (Figure S4E, upper portion). Furthermore, based on published islet epigenome data (Lu et al., 2018), TSS specifically marked with H3K27me3 in β_LO_ cells were highly enriched for poorly transcribed, ‘bivalent’ domains, marked with both H3K27me3 and H3K4me3 (Lu et al., 2018), including the *Cd24a* locus (Figure 4E-G and Figures S4E-G). These data indicate that with increased cell-type resolution, bivalent domains in bulk β-cells largely resolve into subtype-specific active or silent states.

**Figure 4.**
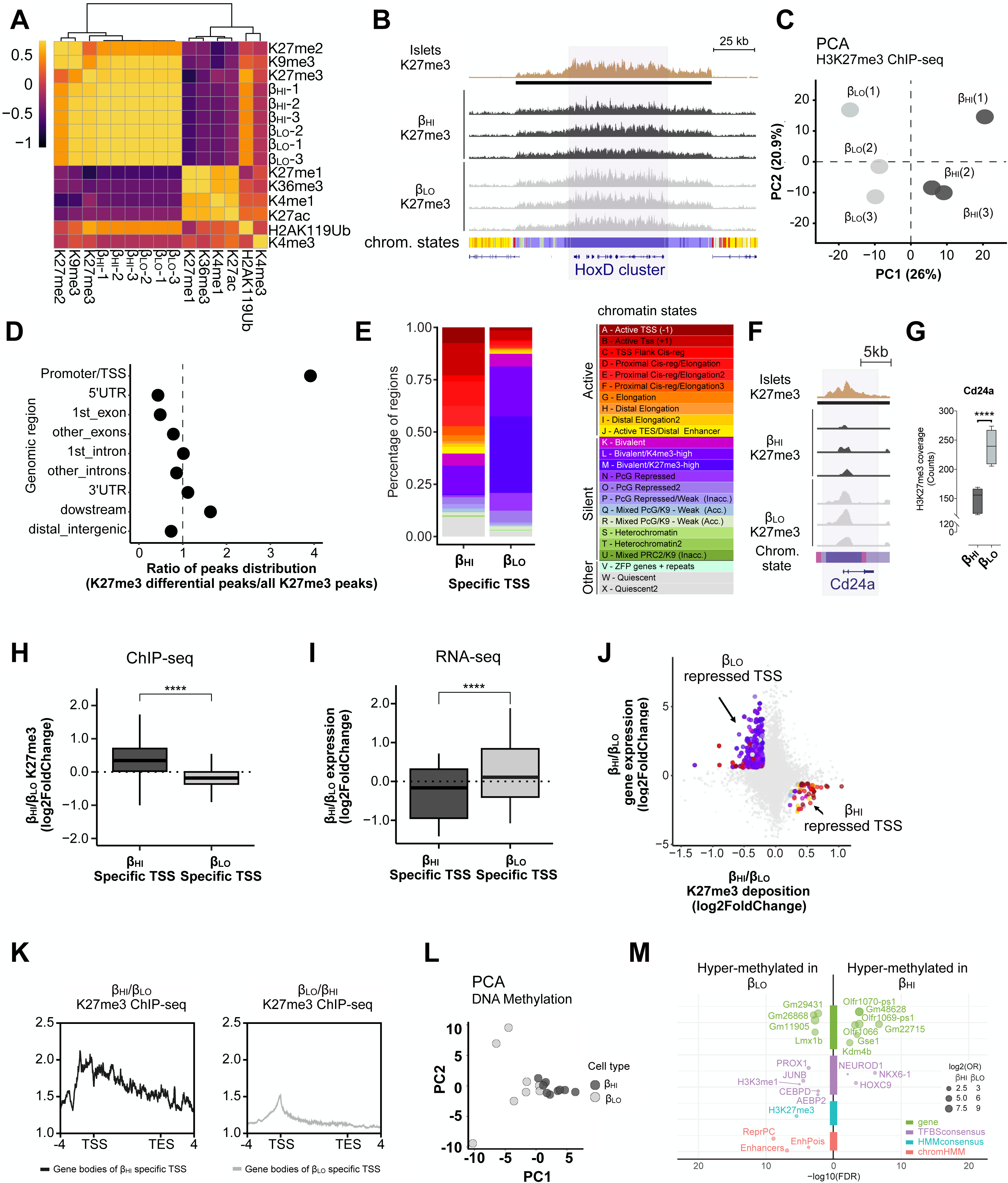
β_HI_ and β_LO_ cells exhibit distinct epigenomes. A. Heatmap showing Spearman correlations of ChIP-seq signals of the indicated histone marks from whole islets, compared to H3K27me3 signals from triplicate experiments of β_HI_ and β_LO_ β-cells. B. Genomic snapshots showing H3K27me3 ChIP-seq tracks from whole islets and purified β_HI_ and β_LO_ cells, as indicated. The HoxD cluster of genes is represented. Horizontal black bars represent H3K27me3 covered broad regions. Colored horizontal bars represent chromatin states, as previously described (Lu et al., 2018) and reproduced in panel (E). C. PCA of H3K27me3 ChIP-seq signals over all identified H3K27me3 peaks, showing reproducible separation of β_HI_ and β_LO_ β-cells. D. Genomic regions’ enrichment among H3K27me3 differential peaks between β_HI_ and β_LO_ β-cells. The dot-plot shows a specific enrichment on transcription start sites (TSS) for H3K27me differential peaks. The distribution of annotated genomic regions over H3K27me3 differential peaks was compared to the same distribution of all identified peaks and plotted as a ratio of percentages (i.e. values >1 mean relative enrichment of H3K27me3 differential peaks over the overall peaks’ distribution, while values <1 mean relative depletion). E. Chromatin states distribution on β_HI_ (left) and β_LO_ (right) H3K27me3-enriched TSS; relative gain of H3K27me3 on active genes (red hues) and relative loss on bivalent genes (purple hues), characterize β_LO_ β-cells. Color-code for chromatin states as previously described (Lu et al., 2018) is reported here F. Genomic snapshots showing H3K27me3 ChIP-seq tracks from whole islets and purified β_HI_ and β_LO_ cells, as indicated. The Cd24a gene is represented. Horizontal black bars represent H3K27me3 covered broad regions. Colored horizontal bars represent chromatin states, as previously described (Lu et al., 2018) and reproduced in panel (E). G. Box plot representation of the Cd24a gene coverage in β_HI_ and β_LO_ cells H. Boxplot showing the ratio of the normalized K27me3 ChIP-seq signal between β_HI_ and β_LO_ cells, on β_HI_ (left, dark-grey) and β_LO_ (right, light-grey) K27me3-enriched TSS. **** = *p*-value < 0.0001, as assessed by *t*-test. I. Boxplot showing the ratio of the normalized RNA-seq signal between β_HI_ and β_LO_ β-cells, on β_HI_ (left, dark-grey) and β_LO_ (right, light-grey) K27me3-enriched TSS. The transcriptional regulation is in line with the reciprocal K27me3 enrichment in panel G. **** = *p*-value < 0.0001, as assessed by *t*-test. J. Scatter plot showing the correlation between β_HI_ / β_LO_ gene expression and H3K27me3 ChIP-signal. Only β_HI_ vs β_LO_ -specific TSS are colored by their chromatin states. K. β_HI_ (left) and β_LO_ (right) β-cells H3K27me3 ChIP-seq signal over the gene bodies of related β_HI_ and β_LO_ -specific TSS’s. The signals are from merged triplicate experiments, and visualized as gene bodies +/− 4 Kb. The coverage profiles show a reciprocal enrichment/depauperation of the K27me3 signal on TSS vs the gene bodies, in β_LO_ and β_HI_ cells, respectively. L. PCA of DNA methylation array signals, showing reproducible separation of β_HI_ and β_LO_ β-cells. M. Enrichment analysis of Differentially Methylated Loci (DMLs) between β_HI_ and β_LO_ within the indicated dataset.

Interestingly, H3K4me3 ChIP-seq performed independently showed that, although differentially marked by H3K27me3, β_LO_-specific TSS’s, including of *Cd24a*, were equally marked with H3K4me3 in both cell types (Figure S4H, I). These data suggest the loci are poised for transcription in β_LO_ β-cells, but active in β_HI_. In line with these findings, differential H3K27me3 deposition at TSSs correlated inversely with transcription at those same genes (Figures 4H-J). These data indicate that the hallmark quantitative differences in H3K27me3 are at least partial drivers of β_HI_ vs β_LO_ differential transcription. Consistent with their higher H3K27me3 staining (Figures 1E-G; -HI), β_HI_ cell specific H3K27me3 deposition was much broader, extending from the TSS well into the gene body when compared to β_LO_-specific enrichments (Figures 4K, S4E). It is also important to remember that coding regions (including the TSS) represent only ∼ 2% of the mammalian genome and that H3K27me3 is primarily found *outside* genic promoter regions (Figure S4D).

Finally, to evaluate whether silent epigenome differences were restricted to H3K27me3, or whether they extended to other cell-type defining drivers of repression, we purified fresh β_HI_ and β_LO_ samples and subjected them to quantitative DNA methylation profiling using Infinium Mouse Methylation BeadChips. Importantly, β_HI_ and β_LO_ cells separated on the first principle component of a methylome PCA (Figure 4L). Enrichment analyses revealed most striking differential DNA-methylation at CpG dinucleotides; almost uniquely at enhancers and H3K27me3 annotated genomic regions (Figure 4M, Figure S4J); at coding regions of β_HI_ cell Lmx1b as well as three annotated pseudogenes; and, at regions with motif enrichments for JUNB, AEBP2, CEBPD, MAFB, ATF3, H3K3me1 and interestingly, the developmental regulator PROX1 (Figure 4M). PROX1 is interesting because the PROX1 locus in humans harbors a genome-wide significant Type-2 diabetes variant (rs340874). Motif enrichments in regions hypermethylated in β_LO_ cells interestingly included those for Nkx6.1 and NeuroD1 (Figure 4M). These findings may suggest a more complete decommissioning of Nkx6.1 and NeuroD1 -associated plasticity in β_LO_ cells. Thus, β_HI_ and β_LO_ cells exhibit distinct H3K27me3 and DNA-methylation control enriched at gene regulatory regions.

### β_HI_ and β_LO_ cells are stably and functionally distinct

Examining the transcriptomic data more carefully, we identified highly co-regulated set of transcripts that were upregulated in β_HI_ cells and specifically transcribed from the mitochondrial - as opposed to nuclear - genome (Figure 5A, green). These differences, importantly, were uncoupled from the regulation of nuclear-encoded mitochondrial genes in all of our data sets (Figure 5A orange, Figures S3D S5A-D) suggesting increased mitochondrial mass in β_HI_ cells. Indeed, PCR-based quantification revealed a near 2-fold increase in β_HI_ cell mitochondrial DNA content (Figure 5B). FACS-based quantification revealed a consistent increase of the mitochondrial protein TOM20 (Figure 5C), and, single cell resolution measurements that indicated consistent and uniform differences across the entire β_HI_ and β_LO_ compartments, with exceptions of a small subset of β_HI_ cells (Figure 5C, histograms). Examination of mitochondrial structure by high-resolution confocal imaging showed that β_LO_ cells had smaller and rounder mitochondria (Figure 5D, E). Finally, TMRM fluorescence, an indicator of mitochondrial activity (Creed and McKenzie, 2019), was increased in β_HI_ β-cells (Figure 5F). These data demonstrate an increased active mitochondrial mass in β_HI_ cells. Thus, β_HI_ and β_LO_ cells exhibit distinct mitochondrial mass, transcription, structural dynamics, and TMRM-associated activity.

**Figure 5.**
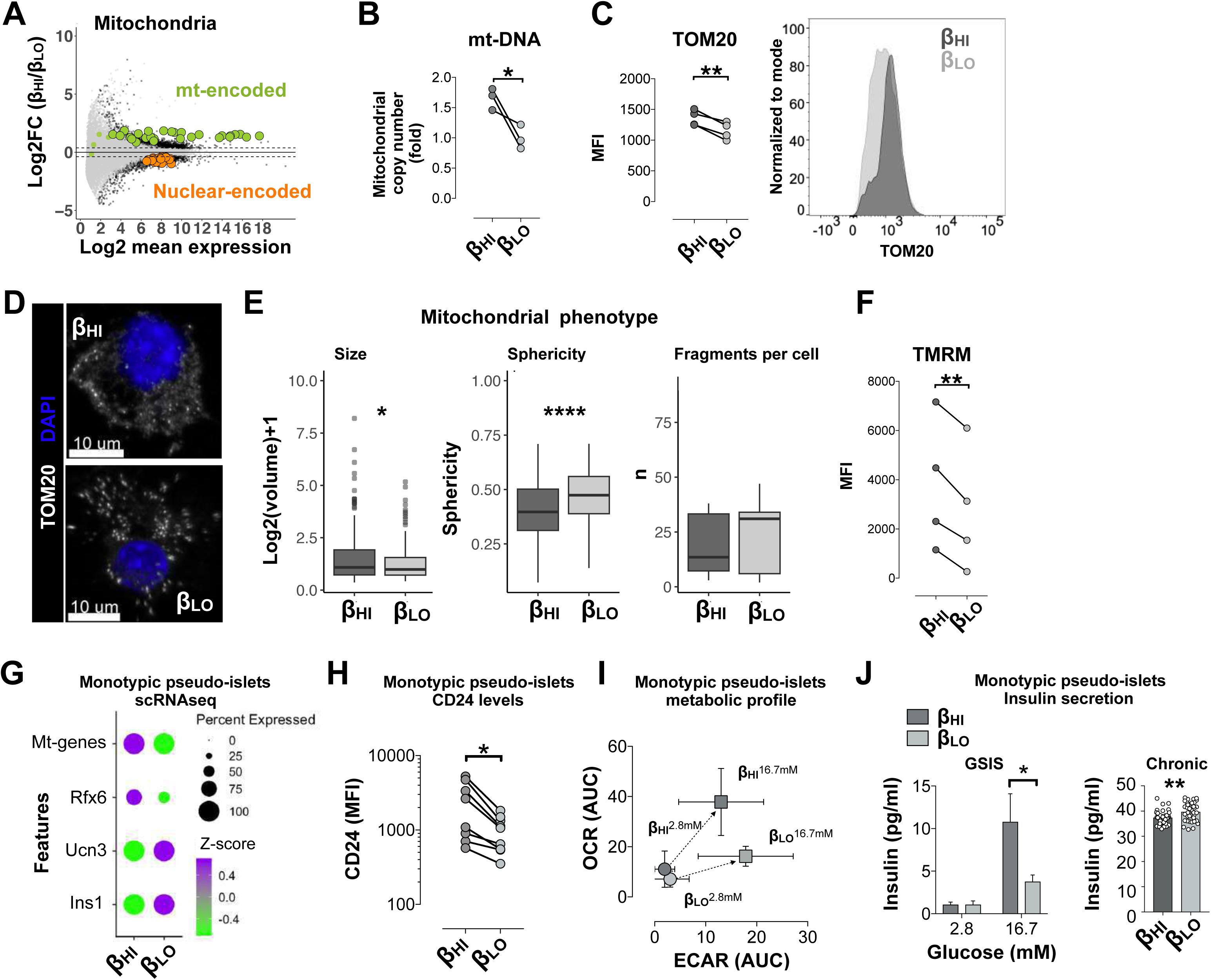
β_HI_ and β_LO_ cells are stably and functionally distinct. A. MA plot showing the fold change in expression generated by comparing β_HI_ over β_LO_ β-cells. Green dots represent mt-encoded mitochondrial genes, orange dots represent nuclear encoded mitochondrial genes listed in S3F. Differentially expressed genes are surrounded by black borders; (*p*-value adjusted for multiple testing < 0.05, with fold change cutoff of 1.33). B. Fold increase in mitochondrial DNA content (copy number normalized to genomic DNA, as measured by qPCR). Each dot represents an independent experiment, n=3. *= unpaired t-test, *p*-value<0.05 C. Dot plot representation of the MFI of TOM20 in the β-cell types, A Representative flow cytometer histogram of TOM20 labeling in the β-cell types. The connected dots represent cells from n=4 individual mice. **= paired t-test, *p*-value<0.01. D. Representative images of TOM20 antibody labeling of one β_HI_ and one β_LO_ cells. fixed β-Cells were first sorted according to their insulin, H3K27me3, and CD24 levels and then labelled with antibody against TOM20 (white) and analyzed at high resolution confocal microscopy. DAPI (blue) was used as counter staining. E. Box plot representations of mitochondrial size, sphericity, and number of fragments per cell. 16 cells were analyzed from n=3 independent mice. *= paired t-test, *p*-value<0.05, ****= paired t-test, *p*-value<0.0001. F. Mean fluorescent intensities (MFI) of TMRM in the β-cell types, connected dots represent cells from n=4 individual mice. **= paired t-test, *p<*0.01. G. Dot plot representation of gene expression levels (z-scored) from scRNAseq of dissociated monotypic β_LO_ or β_HI_ pseudo-islets after 7 days in culture. H. Dot plot representation of FACS measurements of CD24 protein levels in single cells from monotypic β_HI_ or β_LO_ pseudo-islets after 7 days in culture. I. Single spheroid metabolic profiling via Seahorse extracellular flux analysis in basal glucose (2.8mM) and glucose stimulated (16.7mM) conditions. Oxygen consumption rate (OCR) extracellular acidification rate (ECAR) Area under the curves (AUC) are shown in Figure S5I. J. Glucose stimulated insulin secretion (GSIS) and 48 hours, chronic, insulin secretion in single pseudo-islets generated by aggregating 2000 of β_HI_ or of β_LO_ cells. Insulin levels were measured for one hour before stimulation (2.8mM glucose), followed by another hour after stimulation (16.7mM glucose). 25-40 single spheroids were analyzed from n=5 independent experiments. *= two-way ANOVA with multiple comparison correction, *p*-value<0.05.

Mitochondria are a defining regulatory node for β-cell stimulus secretion To test for differences in mitochondrial and secretory function, we therefore FACS-purified β_HI_ and β_LO_ cells and reaggregated them into uniquely β_HI_ or uniquely β_LO_ cell-specific spheroids (monotypic pseudo-islets; Figure S5E). Both readily formed spheroids with no differences in spheroid-forming capacity, rates (Figure S5F) or connexin 36 gap junction gene expression levels (*Gjd2;* Figure S5G). Importantly, even after 7 days in culture, signature mRNA differences remained stable and true to the respective cell-type-of-origin. Specifically, *Ins1* and *Ucn3* were up in β_LO_ monotypic islets, while mitochondrial and *Rfx6* transcripts, as well as surface CD24 were up in β_HI_ cells (Figure 5G, H). β_HI_ and β_LO_ cells therefore maintain their distinctions through dissociation, reaggregation and long-term culture.

Having established monotypic pseudoislets, we performed single spheroid metabolic profiling via Seahorse extracellular flux analysis under basal and glucose-stimulated conditions. In keeping with their increased mitochondria size, sphericity, and membrane activity, β_HI_-monotypic spheroids showed an overall increase in oxygen consumption rate (OCR) relative to extracellular acidification (ECAR) (Figure S5H). Upon glucose stimulation, whereas both spheroid types showed significant ECAR responses, β_HI_-monotypic spheroid alone responded with a substantial OCR response (Figure 5I, Figure S5I). β_HI_ cells therefore are more oxidatively competent in both basal and glucose-stimulated contexts. Finally, we measured GSIS in a parallel single monotypic spheroid setup (Figure S5J). β_HI_ and β_LO_ spheroids both showed robust GSIS (Figure 5K, left panel). Importantly though, β_HI_ spheroids exhibited reproducibly increased GSIS with a near-doubling of insulin secretion upon high glucose challenge. Also noteworthy, under normal culture conditions β_LO_ cells showed a modest increase in chronic insulin output (Figure 5K right panel). Thus, the epigenetically distinct β_HI_ and β_LO_ cells are characterized by stable differences in mitochondrial activity, function and GSIS.

### H3K27me3 dosage controls overall heterogeneity and β_HI_ / β_LO_ cell ratio

To test if H3K27me3 dosage itself is necessary for β-cell sub-type specification and maintenance, we generated animals with a β-cell-specific loss of *Eed* (β-EedKO mice: *Ins1*-Cre^+/-^; *Eed^fl/fl^*). Eed is a critical core subunit of the PRC2 (Polycomb Repressive Complex 2) complex that is responsible for H3K27me3 deposition (Xie et al., 2014). We previously showed that β-cells in β-EedKO mice lose all detectable H3K27me3 between 2 and 8 weeks of age (Lu et al., 2018). Despite the complete loss of β-cell H3K27me3, β-EedKO animals remain glucose tolerant until ∼4 months of age before exhibiting stark and progressive loss of β-cell identity between 4-6 months of age (Lu et al., 2018). To determine if H3K27me3 is necessary for β_HI_ and β_LO_ subtype specification, we isolated islets from 2-month-old β-EedKO animals (immediately after H3K27me3 loss but ∼2 months prior to loss of identity) and performed *SCAN-*seq. Consistent with our previous work (Lu et al., 2018), β-EedKO β-cells at this time-point were devoid of H3K27me3, exhibited normal expression of all key β-cell markers and had normal insulin levels (Figures S6A-C). To determine the effect of H3K27me3 loss on heterogeneity, we performed clustering analysis on cells from both wild-type (Control; Ins1-cre^+^ littermates) and β-EedKO animals. Wild-type β-cells, including β_HI_ and β_LO_ cells, served as our reference (Figure 6A, C and Figure S6D). EedKO β-cells partially overlapped the wild-type transcriptomic space and, additionally, built a trajectory that originated at the wild-type space and formed a trajectory ending in a final relatively tight cluster (Figures 6B-C). Using within cluster *sum of the squared errors* (SSE) we found that although expressing high levels of insulin transcripts and protein EedKO β-cells progressively lose heterogeneity as they move away from the wild-type space, ultimately ‘collapsing’ to a state of low cell-to-cell dispersion (Figures 6B, C and Figure S6B, C), lower even than that observed in either β_HI_ or β_LO_ β-cells alone (Figure 6C and Figure S6E). This conclusion was validated using a dedicated cluster tree analysis that highlighted a substantially lower sub-clustering potential in KO relative to wild-type cells (Figure S6F). These data demonstrate that H3K27me3/PRC2 is necessary *in vivo* for the maintenance of overall β-cell transcriptional heterogeneity, including the separation of β_HI_ and β_LO_ cells.

**Figure 6.**
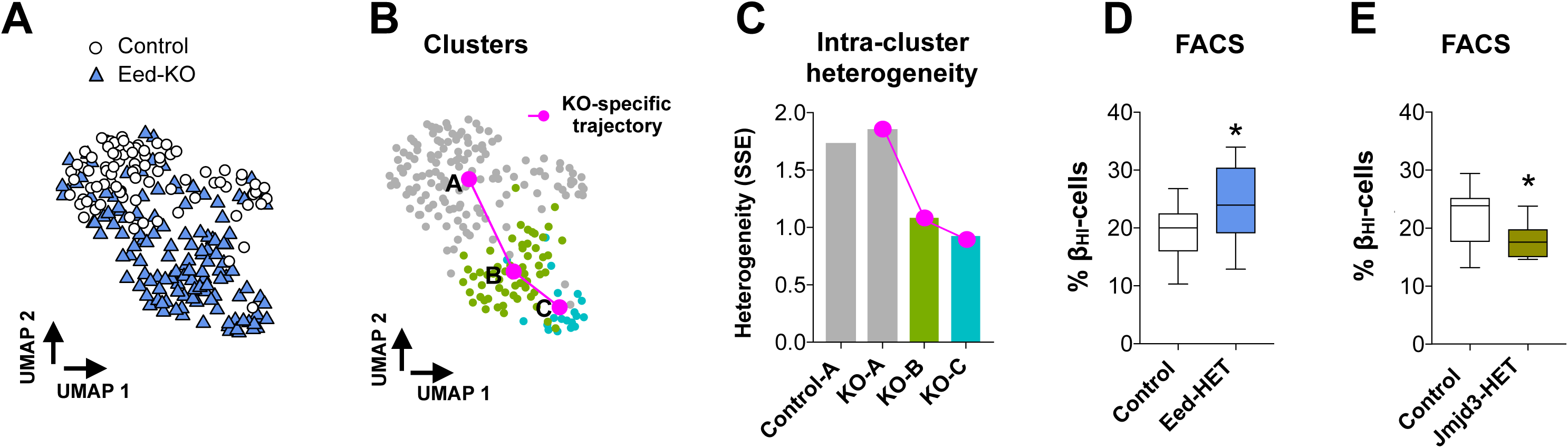
H3K27me3 dosage controls β_HI_ / β_LO_ β-cell ratios and overall heterogeneity. A. UMAP visualization of sorted mouse β-cells that underwent *SCAN*-seq protocol. Colors and shape represent mouse genotypes Eed KO (n= 131 cells) or wild-type (Control; n=83 cells). B. Cluster topology for the data set in (A). Trajectory was inferred by slingshot. Initial clustering was done on all cells, splitting KOs from Controls. KO cluster was further divided into 2 clusters. C. Bar plot showing the intra-cluster sum of squared errors (SSE) per the indicated cluster of cells. As in (B), the magenta line connects the 3 KO groups. D. Box plot representation of the percentage of β_HI_ cells per genotype. Data are medians of Control or Eed-HET mice, n=18 mice each group from 10 or 12 experiments (correspondingly). *= unpaired t-test, *p*-value<0.05. box plots show the median and whiskers indicate min and max values. E. Box plot representation of the percentage of β_HI_ cells per genotype. Data are medians of Control or Jmjd3-HET mice, n=9 mice each group from 6 experiments. *= unpaired t-test, *p*-value<0.05. box plots show the median and whiskers indicate min and max values.

Given that complete PRC2 loss-of-function results in de-differentiation of essentially all β-cells (Lu et al., 2018) (Figure 6B), we sought to assess the consequences of more physiologically-relevant partial H3K27me3 dysregulation. We generated an independent cohort of mice and instead compared β-cells from heterozygous knockouts (β-Eed-Het) and their sex-matched wild-type littermate controls (Ins1-cre^+^; WT) by *SCAN*-seq. The β_HI_ and β_LO_ transcriptomes for each genotype were superimposed (Figures S6G-I). Interestingly however, β-Eed-Het animals showed an increased β_HI_ / β_LO_ cell ratio relative to wild-type (Figure S6J). Since single-cell transcriptomic technologies are neither designed nor intended to provide accurate relative cell counts due to technical confounds, we used FACS-based measures to confirm a reproducible increase in β_HI_ cell numbers in β-Eed-Hets (n=18 mice each group; Figure 6D). Consistent with the heightened GSIS function of Type-1 cells *ex vivo* (Figure 5K), β-Eed-Het animals exhibited both improved glucose tolerance *in vivo* (Figure S6K) and increased insulin secretory function *ex vivo* in isolated islets (Figure S6L). We also examined samples from β-cell specific *Jmjd3* heterozygotes (Ins1-Cre mediated deletion of a conditional *Kdm6b/Jmjd3* allele), a mouse model that *increases* rather than decreases H3K27me3 levels because Jmjd3 is an H3K27me3 demethylase. Indeed, β-*Jmjd3*-heterozygotes showed an equal and opposite cell sub-type distortion (n=9 mice each; Figure 6E). To the best of our knowledge, these represent the first genetic models that trigger β-cell sub-type ratio distortion without impacting cell identity. These data demonstrate that H3K27me3 is a critical determinant of β_HI_ / β_LO_ ratio *in vivo* and (by extension) of the primary axis of β-cell heterogeneity.

### β_HI_ and β_LO_ cells are conserved in humans, and exhibit altered ratios in diabetes

To test whether β_HI_ and β_LO_ β-cells are conserved in humans, we dispersed donor-derived islets provided by the Alberta Diabetes Institute IsletCore and separated them by CD24/H3K27me3 FACS. As in the mouse, human CD24^+^ β-cells were consistently H3K27me3-HI, and CD24^-^ cells were consistently H3K27me3-LO (Figures 7A, B). By confocal imaging, CD24 positive and negative cells were consistently observed within individual islet fragments, validating the presence of both sub-types within individual iselts (Figure 7C). Whereas mouse preparations reproducibly yielded ∼20% β_HI_ cell content (Figure 2D; 19 ± 2 % of all β-cells in young adults), human islet donor preparations exhibited β_HI_ cell numbers ranging from ∼30% to ∼90% of the INS^+^ cell fraction (Figure 7A, Figure S7A). Donor-to-donor variability is well-acknowledged in human islet research (Hart and Powers, 2019) and these data demonstrate a clear need for use of high donor numbers for human islet work and especially when exploring cell-type heterogeneity.

**Figure 7.**
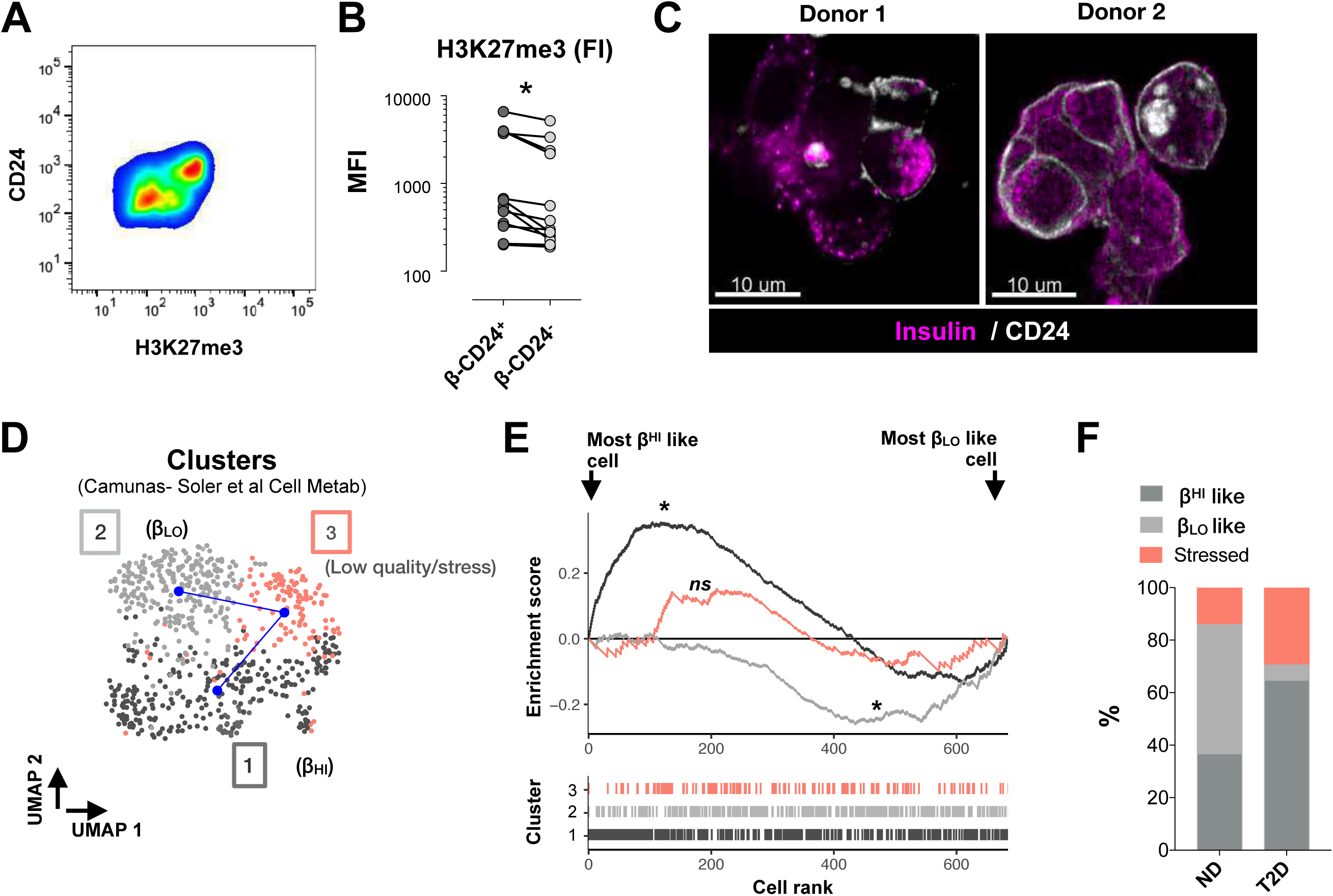
β_HI_ and β_LO_ cells are conserved in humans and their ratio altered in diabetes. A. FACS plot of the fluorescence intensities of CD24 and H3K27me3 in human β-cells isolated from one donor. B. FACS fluorescence intensities of H3K27me3 levels in CD24^+^ compared to CD24^-^ human β-cells, each dot represents the ratio from one donor, n=12 donors. *= paired t-test, *p*-value<0.05. C. Representation of the β-cell surface labeling of CD24 (white) in sub-optimally dispersed, adjacent human islet cells from 2 independent donors. Counter staining of insulin is shown in magenta. D. UMAP representation of the cluster topology of human beta cells. β_HI_/β_LO_ clusters were determined after assessment of expression of the signature, genes reported in Figure 3C. Trajectory was inferred by slingshot. (n=638 β-cells from 11 non-diabetic and 7 T2D donors) E. Custom gene set enrichment analysis (GSEA) representation of β_HI_/β_LO_ signature genes (see Figure 3C). The mean expression (z-score) for the two gene sets was calculated, then the magnitude and direction of differential signatures was determined by calculating the difference in expression between the two gene sets. The cells were then ranked by difference z-score. Plots of cells from all clusters are shown. Cluster 3 had no enrichment. Significant enrichments had *p*-value<0.05. F. Stacked bar plot representation of the percentage of β-cells in each of the clusters shown in (D). Bars split the cluster distributions of non-diabetic (ND) from type 2 diabetic (T2D) donors.

As demonstrated above, these findings validate also through analysis of independent human scRNA-seq/snATAC-seq studies (above), where the major axis of heterogeneity is driven by β_HI_/β_LO_ cell defining H3K27me3 targeted genes (Figures S2H-I). This same axis in humans is distinct from previously reported stress-response associated heterogeneity (Xin et al., 2018) (compare Figure S7B to Figure S2H; UPR= unfolded protein response). To examine potential subtype specific regulation in the context of type-2 diabetes, we analyzed a large single β-cell RNA-seq data which included both diabetic and non-diabetic donors (Camunas-Soler et al., 2020). Interestingly, β-cells grouped into 3 major clusters. One cluster comprised high stress and/or low-quality cells based on gene expression signature and total per-cell transcript counts (Figure 7D, Figures S7C, D). The other 2 major clusters distributed along a β_HI_/β_LO_ β-cell axis according to differentially expressed genes from the two sub-types (Figures S7E, F). Slingshot trajectory analysis demonstrated that both β_LO_ and β_HI_ succumb to stress (Figure 7D). β-cells from T2D donors were enriched in the stressed cluster, confirming previous observations (Shrestha et al., 2021) (Figure 7F, Figure S7E). Importantly, β-cells from T2D donors were also enriched for β_HI_ relative to β_LO_ cells (Figures 7E-F, Figures S7E-F), suggesting a diabetes-specific skew in β_HI_ / β_LO_ cell ratio. Thus, β_HI_ and β_LO_ cells are conserved in humans, and their ratios are affected in T2D.

## Discussion

There is currently no accepted minimal definition for what constitutes a *bona fide* cell type. Historically, stable differences in function, cell-surface protein expression, nuclear and cytological morphology, epigenome configuration, transcriptome and lineage tracing have all been used (independently) to define cell types. And from these collective efforts, we suggest that a key, defining feature of a cell type is *stability* over developmental timescales, and across physiological contexts. Here, we define **‘cell-*state’*** as all forms of heterogeneity - normal or pathological - that can be reflected in any cell type (e.g. differentiation, maturation, aging, circadian rhythm, fasting/feeding/diet, transcriptional bursting). These are ubiquitous in the body and tend to have temporal characteristics. We propose that ‘**cell-*types****’ (or sub-types)* be reserved for cell populations whose distinguishing features **i.** arise normally over developmental time-scales; **ii.** are reproducibly and stably detected across a wide range of contexts (ages, circadian time, diets, disease); **iii.** exhibit wide-spread and stable differences in their active *and silent* epigenomes, transcriptomes, surface protein expression and function; and ideally, **iv.** that maintain these differences through long-term culture under identical conditions. We propose ‘*sub-population’ or ‘subset’* be used where these distinctions are not known or intended. So, while single-cell methods like scRNA-seq are ideally suited for detecting heterogeneity (i.e., differences), they do not necessarily generate the most appropriate data for defining a cell type (or sub-type). Here, we used a combination of bulk and single-cell technologies, cell sub-type specific spheroid culture, imaging, mitochondrial and functional analysis to identify new epigenetic axis that defines two primary β-cell sub-types (βHI and βLO cells) that are distinct by 7 sets of criteria (function, FACS markers, epigenome configuration, transcriptome, nuclear and cytosolic ultrastructure, and morphology).

### Literature context

One of the major challenges facing the field has been integration of heterogeneity datasets with insights made using disparate technologies and occasionally with unidimensional data. These challenges stem partly from technological inadequacies of scRNA-seq itself, and from rational use of genetic tools such as transgenic reporters that are by definition artificial, generate heterogeneity themselves and may not express reproducibly in all cells of a given compartment (e.g. widely used Nkx6.1 reporter lines while highly valuable express in ∼80% of β-cells (Liu et al., 2021)). We focused our approach on evaluating heterogeneity across *all* β-cells. We did this by leveraging insulin antibody positivity and evaluated heterogeneity initially using FACS that scales effectively over 5 orders of magnitude. For these reasons, we conclude that β_HI_ and β_LO_ cells (at least in mice) comprise >90% of the adult β-cell compartment. Outside these ∼90%, we identify a CD24^high^ SST/INS double-positive sub-population that deserves careful exploration considering recent work highlighting transdifferentiation and islet endocrine cell plasticity (Bramswig et al., 2013; Chera et al., 2014; Thorel et al., 2010; van der Meulen et al., 2017).

Notably absent from our single-cell data are high-UPR clusters. In our hands, the immediate transcriptome ‘freeze’ enabled by the immediate fixation step of *SCAN-*seq eliminates high-UPR cells that we find in parallel tests from the very same islet isolation batch where the cells are simply processed without fixation (ie. CELseq2 alone; not shown). These data suggest strongly that a substantial fraction of the stress-response signature observed in single-cell genomic data likely results from isolation, fluidics and sorting steps. These ideas are consistent with recent systematic examinations of such confounds (Marsh et al., 2022; Nguyen et al., 2018). Given the importance of UPR in β-cell biology (Engin et al., 2013; Lee et al., 2020; Shrestha et al., 2021; Xin et al., 2018), the experimental design for examining this process appears to be especially critical.

Populations of early responding and highly interconnected ‘hub’ β-cells (1-10% of total) classified by in situ Ca^+2^-responsiveness, have been reported with suggested mitochondrial functional heterogeneity. Those cells exhibited low insulin protein, signatures of immaturity, and increased metabolic function without a difference in TOM20 (Johnston et al 2016). A related set of ‘leader’ cells reported by the same group exhibit transcriptional enrichment of chromatin regulators (Chabosseau et al bioRxiv, 2022), and, appear to derive almost entirely from the smaller of two major clusters of pancreatic β-cells by scRNAseq (Salem et al 2019). Tandem single-cell resolution electrophysiological and RNAseq profiling by PATCH-seq similarly identified gradients of electrophysiological responsiveness associated with physically smaller, *Rbp4-*enriched, β-cells that showed lower exocytosis under chronic conditions. Our functional analyses and examination of transcriptional patterns of these factors (Figure S2K, L) suggest that the PATCH-seq *Rbp4*-enriched cells, and potentially also ‘leader cells’, comprise one of two subsets of β_HI_ cells.

Our findings demonstrate that the major axis of β-cell heterogeneity is formed at least in part by epigenetic silencing machinery. Interestingly, several genes coding for reported markers of β-cell heterogeneity (including *Cfap126*, *Rbp4*, and *Ffar4*) as well as a large proportion of β-cell *disallowed* genes (Pullen et al., 2010) map to H3K27me3-marked regions. Our examinations indicate that the axes of maturation, which are marked for instance by Cfap126, Mafa, and CD81 (Bader et al., 2016; Nasteska et al., 2021; Salinno et al., 2021), exist *within* both β_HI_ and β_LO_ cell compartments suggesting that maturation gradients exist *in vivo* for both cell-types.

Along those same lines, an immediate question of interest is precisely how β_HI_ and β_LO_ cells are specified, and how their maturation and maintenance couples to metabolic demands. We find both subtypes are present from early to late life in mice, they both proliferate, they are stable for at least 7 days under identical culture conditions, and they are metabolically quite distinct. mTOR and AMPK signaling play crucial roles in β-cell maturation process (Helman et al., 2020; Jaafar et al., 2019) and interestingly β-cell specific mTOR deficient mice have lower levels H3K27me3 and upregulation of a group of ‘immature’ genes (Ni et al., 2022). Interestingly, in transcriptome data we observe reciprocal mTOR regulation in β_HI_ and β_LO_ cells (increased expression of negative regulators of mTOR Tsc1, Tsc2, Ubr1, and Rictor in β_HI_; upregulation of the positive regulators Lamtor2, Golph3, Rheb, Deptor, Lamtor1 in β_LO’s;_ not shown). This observation suggests that fidelity of cell sub-type identity may be continuously reinforced by regulation of TOR signaling.

Finally, relative to work that certainly motivated the field by identifying sortable clusters of β-cells based on surface antigen FACS analysis (Dorrell et al., 2016) our findings are inconclusive. Assessment of their differentially expressed genes in our SCAN-seq data set highlighted differences in the expression of their β1/β2 specific genes (Figure S2K, M UMAPs panel). Similar to our findings (Figure 3F), Rfx6 and Mafb appeared as a key transcription factors differentiating their ST8SIA1^−^ β1/β2 cells. RFX6 also had promoter accessibility in INS^high^ state cells observed elsewhere (Chiou et al., 2021). The data suggests therefore that those may be β_HI_ cells though Rfx6 also aligns with maturation gradients within both β_HI_ and β_LO_ cell clusters by scRNAseq.

### Next steps

Next important steps for the field will be to test whether β_HI_ and β_LO_ cell differences can be harnessed for stem-cell based islet replacement strategies, and to test whether β_HI_ or β_LO_ cells are preferentially dysregulated in classically-defined (T1D, T2D, MODY, gestational diabetes, etc.) as well as newly emerging diabetes sub-types (Ahlqvist et al., 2020) (SAID, SIDD, SIRD, MOD, MARD). Key steps towards these ends will be to identify additional surface antigens for multi-marker separation that are robust against experimentally induced and donor-to-donor variation. An additional priority will be to understand the up- and down-stream factors that drive sub-type specification and the mechanisms that link Eed and Jmjd3 dosage to β_HI_ / β_LO_ ratio control.

Indeed, to the best of our knowledge, the β-Eed-Het and β-Jmjd3-Het animals represent the first examples of reciprocal genetic models that specifically skew β-cell subtype ratios (heterogeneity) *in vivo.* In the case of the β-Eed-Het animals, scRNAseq data indicate that this happens without impacting transcriptional identity of either β-cell subtype indicating ratio control is independently regulated from identity. The data suggest that even subtle changes in the H3K27me3 levels, such as the reduction observed in T2D (Lu et al 2018) have the potential to modulate β_HI_/β_LO_ cell ratios over the long-term especially in human disease timeframes. Notably, the observed increase in β_HI_/β_LO_ cell ratio in T2D (Figure 7F) is consistent with a model where H3K27me3 dysregulation causes skewing of cell subtype ratio in T2D. Along similar lines, the heightened gluco-regulatory phenotype observed in β-Eed-Het animals indicates that manipulation of β-cell ratios could constitute a desirable therapeutic goal in the context of metabolic disorders, and it suggests that the skewing observed in T2D constitutes a form of compensation. The data suggest that *low-dose* or *intermittent* Ezh1/2 or Eed inhibition could serve a role in improving those methods aimed at generating β-cell replacements from stem or iPS cells. Substantial activity in epigenetic inhibitor space has already identified a range of *in vivo* tolerated compounds with specificities for PRC2 catalytic and other subunits.

### Concluding thoughts

Finally, our analysis, which leverages the respective powers of scRNAseq (for characterizing cell-to-cell heterogeneity), bulk RNAseq (for deep and accurate transcript quantification), and monotypic spheroids (for functional analysis), as well as our independent analysis of the 6 publicly available datasets in Figure S2G-I, do not necessarily support the prevailing view that there exists a *common immature* β-cell sub-*<u>type</u>.* Rather, they support the notion that cell-state gradients (of maturity, cell cycle, aging, disease, circadian and diet, etc) exist across each of 2 or more highly specialized β-cell subtypes. β_HI_ and β_LO_ cells both exhibit robust and equal expression of essentially all known terminal differentiation markers despite clear maturation gradients readily detectable across each. These nuances are important as the community works towards a common framework for β-cell heterogeneity.

## Acknowledgements

We would like to thank Prof. Thomas Jenuwein for stimulating interactions and enduring research support. We would like to acknowledge excellent technical support of the MPI-IE and VAI core services and in particular Laura Arrigoni, Chiara Bella, Leila Rabanni. We thank Dr Sherri L Christian for providing the CD24 knockout mice. Human islets for research were provided by the Alberta Diabetes Institute IsletCore (www.bcell.org/adi-isletcore) and the Clinical Islet Laboratory at the University of Alberta in Edmonton with the assistance of the Human Organ Procurement and Exchange (HOPE) program, Trillium Gift of Life Network (TGLN), and other Canadian organ procurement organizations. Islet isolation was approved by the Human Research Ethics Board at the University of Alberta (Pro00013094). All donors’ families gave informed consent for the use of pancreatic tissue in research. We thank Dr Darrell Chandler for critical evaluation of the manuscript. FUNDING: ERC, Van Andel Institute, TR01, R21, EFSD.

## Author contribution

ED and JAP conceived the project. ED designed, performed, and analyzed all experiments unless stated differently. LF analyzed the ChIP-seq and RELACS datasets, and together with SA the DNA methylation array datasets. VW helped perform the *in vivo* experiments and islet isolations. SH supported the bioinformatics work and code for the *SCAN*-seq multimodal pipeline. LF, SA, BJ, AS, PS, DS provided support for the bioinformatic analysis. KDH, IP, VK, RC, TTL, AL, BG, TG, and AI helped perform experiments. JAP supported study design, data analysis, and acquired financial support. ED and JAP wrote the original draft. All co-authors reviewed and edited the manuscript.

## Methods

### Animal Husbandry

All animals were maintained on a normal chow diet with 15% fat (Ssniff GmbH), fed *ad libitum* with free access to water (HCl acidified, pH 2.5-3) under controlled humidity and temperature with a 12-hour light and 12-hour dark cycle. High fat diet fed mice were fed with 60% kcal% fat diet (Research Diet) for 3 days or 4 weeks. All animal studies were performed with the approval of the local authorities in either in Germany (Regierungspräsidium Freiburg, Germany) under license number 35-9185.81/G-16/120 or approved by the Institutional Animal Care and Use Committee at the Van Andel Research Institute, Grand Rapids, MI, USA under the animal use protocol number 21-08-023.

### Genetically modified Mice

The CD24 knockout(Nielsen et al., 1997) mice were kindly provided by Sherri L Christian. Breeding pairs of Ins1-cre (Thorens et al., 2015) (B6(Cg)-Ins1tm1.1(cre)Thor/J were purchased from Jackson laboratories). Eed^fl/fl^, Kdm6b^fl/fl^, and YFP-reporter (B6.129X1-Gt(ROSA) 26Sortm1(EYFP)Cos/J) transgenic mouse line (C57B6/J) were kindly provided by Stuart Orkin, and Thomas Boehm, respectively. To generate β-cell reporter mice with Eed deficiency, Eed-floxed animals were crossed with YFP harboring-Ins1-cre positive animals. All mice had been backcrossed for over 10 generations before any phenotyping was initiated. Experimental mice were all males, unless otherwise stated. Age of the mice used for individual experiments are specified accordingly.

### Islet Isolation

Adult pancreata were perfused through the common bile duct using a 30-gauge needle with Collagenase 4 solution (dissolved in 1x HBSS, 10mM HEPES at a concentration of 1mg per mL), this step was excluded for neonatal islet isolation. Then the pancreata were dissected and incubated 30 minutes in the same collagenase solution. Islets were purified as previously described (Dror et al., 2017). The Isolated islets were hand-picked and cultured in complete media (RMPI-1640 containing 11 mM glucose, 10% FBS, 0.1% Penicillin/Streptomycin, gentamicin and Amphotericin B; Thermo Fisher) and maintained at 37°C in 5% CO_2_ environment to allow their recovery.

### Human pancreatic islets

Human islets were from and the Alberta Diabetes Institute IsletCore(Lyon et al., 2016) and the Clinical Islet Laboratory at the University of Alberta, respectively. They were isolated from pancreata of cadaveric organ donors in accordance with the local Institutional Ethical Approvals (Pro00013094). Islets were cultured in CMRL-1066 medium containing 5 mmol/l glucose, 100 units/ml penicillin, 100 μg/ml streptomycin, 2 mM Glutamax and 10% FCS (Invitrogen) in humid environment containing 5% CO_2_.

### Islet dispersion and single cell labeling for FACS

For islet dispersion, islets were incubated in accutase for 4 minutes at 37°C, then gently pipetted using a 1mL pipette for 20 times. Immediately after, single cell suspensions were examined under the microscope, and validated cell suspensions were washed with 2mL of ice cold FACS buffer (PBS, 0.5% BSA, 5mM EDTA) or ice-cold PBS (in case of subsequent fixable viability labeling; 5 minutes zombie dye on ice). Cell surface CD24 (mouse: Thermo Fisher 48-0242-82, human: Biologend, 311122, 1:250), CD45 (Thermo Fisher, 11-0451-82, 1:200), and CD31 (BD pharminfen, 558738 1:200) labeling was done for 30 minutes on ice (diluted in FACS buffer). After washing, cells were fixed in 1% methanol free formaldehyde (Thermo fisher, 28906, 1mL; diluted in RPMI; freshly made) for 15 minutes, the reaction was quenched with glycine (end conc. of 125mM) and cells were washed with additional 1mL of FACS buffer. Intracellular labeling of insulin (sc-8033, 1:100), glucagon (sc-51459, 1:100), somatostatin (sc-55565, 1:100), pancreatic polypeptide (sc-514155, 1:100), MKI67 (47-5698-82, 1:200), TOM20 (Abcam, 1:500), and chromatin labeling using conjugated H3K27me3 (Origene, TA347154 conjugated using mix-n-stain CF555 kit, Sigma), H3K4me3 (C15410003, conjugated using mix-n-stain CF405 kit, Sigma), H3K36me3 (C15410192, conjugated using mix-n-stain CF405 kit, Sigma), H3K9me3 (C15410193, conjugated using mix-n-stain CF488 kit, Sigma), H3K27ac (C15410196, conjugated using mix-n-stain CF488 kit, Sigma) at final concentrations of 5ug/ml, was done in permeabilization buffer (eBioscience, 00-8333-56). For data in Figure 1A, H3K27me3 was labeled together with either H3K9me3 and H3K36me3 or H3K27ac and H3K4me3. The presented data in Figure S1F shows additional labeling with H3K27me3 (C15410195, Diagenode). Unless stated differently-insulin positive β-cells were analyzed and sorted while excluding glucagon, somatostatin, pancreatic polypeptide, CD31, and CD45 positive cells that were included in a ‘dump channel’; 488, which contained all of the antibodies that are specific for the unwanted cells. Washing between steps was determined differently; live cells were centrifuged at 190g, fixed cells at 350g and fixed-permeabilized cells were centrifuged at 500g, all for 4min at 4 degrees. For experiments with subsequent extraction of RNA, all the steps were done in the presence of RNase inhibitor (recombinant RNasein, Promega, N2511) diluted 1:4,000 for washing steps or 1:400 for incubation while staining and for buffers in the tubes containing the sorted cells followed by snap freeze and storage at –80°C.

### *SCAN*-seq

The new multi-modal ‘single cell *S*urface, *C*ytoplasmic *A*nd *N*uclear staining and RNA sequencing (*SCAN*-seq) of FACS labeled and fixed single β-cells (described above) was built off of the CEL-Seq2 method(Hashimshony et al., 2016; Lu et al., 2018). Insulin positive cells were index-sorted into 384 well plates containing 384 unique barcodes (supplemental table), Single cells were sorted in 384-well plates (Bio-Rad Laboratories, HSP3801) containing lysis buffer and mineral oil (Sigma, M8410) using BD FACS Aria FUSION. The sorter was run on single-cell sort mode with index sorting. Doublets were excluded using pulse geometry gates (FSC-W × FSC-H and SSC-W × SSC-H). Importantly, cells from all conditions/biological replicates were equally distributed into wells of all sorted plates from the same experiment to enable optimal batch correction in case of evident plate bias in the protein of transcriptional data. After the completion of sorting, the plates were centrifuged for 2 minutes at 2,200 g at 4°C, snap-frozen in liquid nitrogen and stored at -80C for up to two weeks until processed. 160 nL of reverse transcription reaction mix and 2.2 mL of second strand reaction mix was used to convert RNA into cDNA. cDNA from 384-cells was pooled together before the clean-up and *in vitro* transcription, generating one library from one 384-well plate. 0.8 mL of AMPure/ RNAClean XP beads (Beckman Coulter GmbH, Germany) per 1 mL of sample were used during all the purification steps including library cleanup. Libraries were sequenced on a single lane (pair-end multiplexing run, 100 bp read length) of an Illumina HiSeq system targeting 200,000 reads per cell.

### Bulk-cell RNA-seq

Total RNA from 1,000 H3K27me3 HI/LO β-cells or from 50,000 sorted β_HI_ and β_LO_ cells was extracted using the miRNeasy FFPE Kit (QIAGEN, 217504), followed by the NEBNext® Single Cell/Low Input RNA Library Prep Kit for Illumina® (E6420L). Library fragments of 350 ± 20 bp were obtained, and the quality was assessed using a Fragment Analyzer (Advanced Analytical). Barcoded libraries were subjected to 70bp pair-end sequencing on the Illumina HiSeq 2000.

### ChIP-seq

Chromatin from snap-frozen pellets of the sorted β_HI_ and β_LO_ cells (3 biological replicates each been prepared using the NEXSON procedure(Arrigoni et al., 2016) to a 100-800 bp fragment size distribution. Sheared chromatin was controlled for size distribution and cell number. For this, 2 ml of 20 mg/ml Proteinase K was added to a small chromatin aliquot (5 ml) of each sample. Volumes were adjusted to 20 ml using buffer EB (Qiagen). Samples were then reverse-crosslinked by incubating them at 50 °C for 30 min, followed by incubation at 65 °C for 30 min. DNA concentration was measured using Qubit dsDNA HS assay to estimate cell concentration (one mouse diploid cell contains approx. 6.6 pg of DNA). Samples were then purified using Qiagen MinElute columns, and DNA fragment size distribution was checked by capillary electrophoresis (Agilent Fragment Analyzer).

Before ChIP all chromatins were normalized to the same cell number using shearing buffer. Normalized chromatins have been diluted 1:2 in 1X Buffer iC1 (supplemented with protease inhibitor cocktail) from the Diagenode iDeal ChIP-seq kit for histones (C01010173). Each chromatin sample containing about 6,000 cells was incubated with 1 µg of anti-H3K27me3 antibody (Diagenode, C15410195, lot. A1811-001P). ChIP was performed using the automated platform SX-8G IP-Star platform (Diagenode) under the program “ChIP indirect method”. The antibody-chromatin incubation lasted 10h, followed by 3 hours of bead incubation (protein-A conjugated), and 5-minutes beads washes. Ten percent of the original chromatin was used as input. After the DNA elution, ChIP and input samples were de-crosslinked and purified using the Qiagen MinElute columns.

Libraries were prepared on an automated liquid handler (Biomek i7) using the NEBNext Ultra II DNA library preparation kit (NEB, E7645), according to the manufacturer’s instructions and without size selection. Libraries were sequenced paired-end on the Illumina NovaSeq platform.

### RELACS

The H3K4me3 ChIP-seq was performed using RELACS as previously described (Arrigoni et al., 2018). Briefly, 50,000 β_HI_ and β_LO_ cells cells were thawed in RELACS lysis buffer (10 mM Tris-HCl [pH 8], 10 mM NaCl, 0.2% Igepal, 1× Protease inhibitor cocktail) and the nuclei were isolated by sonication using the NEXSON procedure(Arrigoni et al., 2016). To digest the chromatin, 25 µl of 10× CutSmart buffer (NEB), 2.5 µl 100× Protease inhibitor cocktail and 1 µl of CviKI-1 (5 U/100,000 nuclei, NEB R0710S) were added. The digestion reaction was incubated overnight at 20 °C. End repair and A-tailing was performed and customized adapters were ligated to the fragments. Once barcoded, the samples were pooled together. Chromatin was then sheared by sonication (Covaris E220, MicroTubes, 5 min, peak power 105, duty factor 2, cycles burst 200). This chromatin was used for automated ChIP (Diagenode, C15410003) with the IP-Star Diagenode system. IPs and Inputs were decrosslinked, DNA was purified and libraries were prepared using the NEB Ultra II DNA Library Prep Kit for Illumina (E7645S and E6440) following the manufacturer’s instructions. Integrity and size-distribution of the samples was assessed before and after library preparation by running on Fragment Analyzer (Advanced Analytical).

### DNA methylation array

Genomic DNA was extracted from fixed and sorted β_HI_ or β_LO_ using the Zymo Research Quick-DNA Microprep Plus Kit (Zymo Research, Irvine, CA USA), according to manifacturer’s instructions. DNA samples were next quantified by Qubit fluorimetry (Life Technologies) and bisulfite converted using the Zymo EZ DNA Methylation Kit (Zymo Research, Irvine, CA USA) following the manufacturer’s protocol, with the specified modifications for the Illumina Infinium Methylation Assay. After conversion, the bisulfite-converted DNA was purified using the Zymo-Spin binding columns and eluted in Tris buffer. Following elution, bisulfite-converted DNA was processed through the Illumina mouse methylation array protocol. The bisulfite-converted DNA samples was first processed using the Infinium HD FFPE DNA Restore kit workflow. To perform the Infinium assay, converted DNA was denatured with NaOH, amplified, and hybridized to the Infinium bead chip. An extension reaction was performed using fluorophore-labeled nucleotides per the manufacturer’s protocol. Array BeadChips were scanned on the Illumina iScan system and signals where assigned by using Illumina Genome Studio v2011.1 software, to produce IDAT files. CpG probe selection (Zhou et al., 2022) included an array of target and random, controls sequences. The DNA methylation array analysis was performed using SeSAMe (Zhou et al., 2018) and its wrapper pipeline SeSAMeStr (10.5281/zenodo.7510575). Nine biological replicates of β_HI_ and β_LO_ were compared. Data pre-processing and quality controls were performed using SeSAMe default parameters and the pre-processing code ‘TQCDPB’. All samples showed a detection rate > 93% and no dye bias. PCA analysis of beta values was performed within the SeSAMeStr pipeline, using the R function ‘prcomp’. In all differential analysis, the effect size cutoff was set to 0.05 (i.e., 5% dffertential DNA methylation) and the p-value cutoff was < 0.05. Further visualization of SeSAMe/SeSAMeStr output data, was perform in R using Rstudio.

### Re-aggregation of islet spheroids

Islets were Isolated as described above from the β-cell reporter mice that were generated by crossing Ins1-Cre mice(Thorens et al., 2015) with the YFP reporter mice B6.129X1- Gt(ROSA)26 Sortm1(EYFP)Cos/J floxed-stop-YFP. After overnight recovery, islets were dispersed as described above to achieve single-cell suspensions for CD24 labeling. Live, CD24^-^ or CD24^+^ YFP^+^ β-cells were sorted into tubes containing 1x HBSS (GIBCO) 0.5% w/v BSA (Serva) and 24mM HEPES (Sigma). Sorted cells were centrifuged 200g for 4 minutes at 4°C degrees and resuspended in mouse islet media (see islet isolation section above) at the concentration of 10 cells/µL then distributed into 96 well plates (U bottom - Nunclon Sphera)- 200µL/well (i.e.2000 cells/well). To determine spheroid formation kinetics, the plates were incubated inside a real-time quantitative cell imaging system (Incucyte®) that was set to image cells as they aggregate every 15 minutes for 3 days.

### Microscopy and Image Quantification

For the analysis of islets, whole, dispersed, or sorted β-cell cells, at least three animals of each condition were analyzed. Samples were stained live or fixed with 1% methanol free formaldehyde, permeabilized with 1x permeabilization buffer (00-833-56 Invitrogen) and stained for the indicated antigens/proteins with sample-type-specific adjustments; single cells were incubated for 30 minutes on ice while whole islets were incubated rotating overnight at 4 degrees. Images were acquired using the LSM880 confocal microscope (ZEISS) with the Airyscan super-resolution (SR) mode turned on. An identical threshold was applied to all images from the same channel to exclude background signals. H3K27me3 positivity and intensity in DAPI positive nuclei as well as TOM20 based analysis of the mitochondrial structure was quantified using Imaris version 9.3.1 in a blinded manner.

### Oral glucose tolerance test

For the oral glucose tolerance test (OGTT), mice were fasted for 6 hours (8:00-14:00), after which basal blood glucose was measured. Mice were given glucose (1 g/kg) by oral gavage. Blood glucose levels were measured using a OneTouch Vita blood glucose meter at 0, 15, 30, 60, and 90 minutes after glucose.

### Measurements of Oxygen consumption rate (OCR) and extracellular acidification rate (ECAR)

An XF96e Extracellular Flux analyzer (Seahorse Biosciences) was used to determine the bioenergetics profile of single monotypic pseudo-islets. Prior to the assay, monotypic pseudo-islets, 2000 cells each were incubated in unbuffered DMEM (Seahorse Biosciences). Then, single spheroids were hand-picked and under the microscope were added into the middle of a 96-spheroid ploy-L-lysine coated microplate (Seahorse biosciences). After two 2-minute basal measurements, glucose was injected into the media (16.7mM end concentration) and the oxygen consumption and extracellular acidification rates were measured for 4 times, 2 minutes wach time. Between every measurement a 5 second mixing step was followed by 5 second waiting step. Wells with readouts lower than background measurements were excluded from further analysis.

### Glucose stimulated insulin secretion

Single re-aggregated cell-type specific pseudoislets (organoids) or overnight recovered whole islets were pre-incubated for 30 minutes in pre-equilibrated Krebs-Ringer bicarbonate buffer (KRB; 115 mM NaCl, 4.7 mM KCl, 2.6 mM CaCl2 2H2O, 1.2 mM KH2PO4, 1.2 mM MgSO4 7H2O, 10 mM HEPES, 0.5% bovine serum albumin, pH 7.4) containing 2.8 mM glucose. Single, β_HI_ or β_LO_ pseudoislets were individually transferred into a V-shaped well of a 96 well plate, each well containing 50µL 2.8mM glucose-KRB. Pseudoislets were incubated at 37°C for 1 hour for basal secretion. Then, individual pseudoislets were collected, washed in PBS and then incubated in 16.7mM glucose KRB- containing V-shaped well for 1 hours.

Whole islets from all sizes were added (5 per well of a 48 well plate), containing either 2.8mM glucose (basal) or 16.7mM glucose (stimulated) Krebs-Ringer buffer and were incubated at 37°C for 1 hour. The islet supernatants were collected after incubation. Each step and kept on ice. Supernatants were centrifuged (2000g 5min 4°C), transferred to new 96 well plate, and stored at −20°C for later insulin measurements using ultrasensitive insulin ELISA (Mercodia).

### Mitochondrial Membrane Potential by FACS Analysis

Ins1-YFP islets were allowed to recover overnight, dissociated as described above, and washed twice with Krebs solution containing 4 mM glucose. For detection of the mitochondrial membrane potential, dissociated islet cells were incubated with 10 nM of the fluorescent probe TMRM (Life Technologies) solution containing 4 mM glucose. Cells were washed with PBS once, scored by FACS using BD Symphony, and analyzed by Flowjo.

### Mitochondrial DNA quantification

Mitochondrial and genomic DNA was isolated from FACS sorted β_HI_ and β_LO_ cells according to commercially available isolation kit and according to the manufacturer instructions. (Absolute Mouse Mitochondrial DNA Copy Number Quantification qPCR Assay Kit (AMMQ) Catalog #M8948)

### Statistical analysis

#### Bulk RNA-seq analysis

RNA-seq was performed with at least three independent biological replicates. Raw sequences from the biological replicates were aligned. Reads for mouse bulk RNA-seq datasets were mapped against mouse genome version mm10 with the snakePipes2 RNA-seq pipeline. Differential expression analysis was performed with DEseq2 (Love et al., 2014). Genes with counts <2 were excluded from differential analysis. Differential genes were called with an FDR threshold of 0.05 and a fold change of 1.33. After QC, and exclusion of lowly expressed genes (>2 counts) differential expression of the raw counts was performed using DESeq2 v1.34.1. Samples were batch-corrected using Limma, and normalized count matrices were inspected using PCA. Gene Set Enrichment Analysis (GSEA) of DE results was performed with fgsea R-package. Enrichment maps were generated in Cytoscape (Shannon et al., 2003). Motif enrichment analysis on β_HI_- specific TSS was perfomed using HOMER v4.11 (Heinz et al., 2010) function ‘findMotifsGenome.pl’, with ‘-size given -mask’ options, and using the transcriptionally unchanged TSS between β_HI_ and β_LO_ cells, as background control.

## Supplemental information

**Figure S1.**
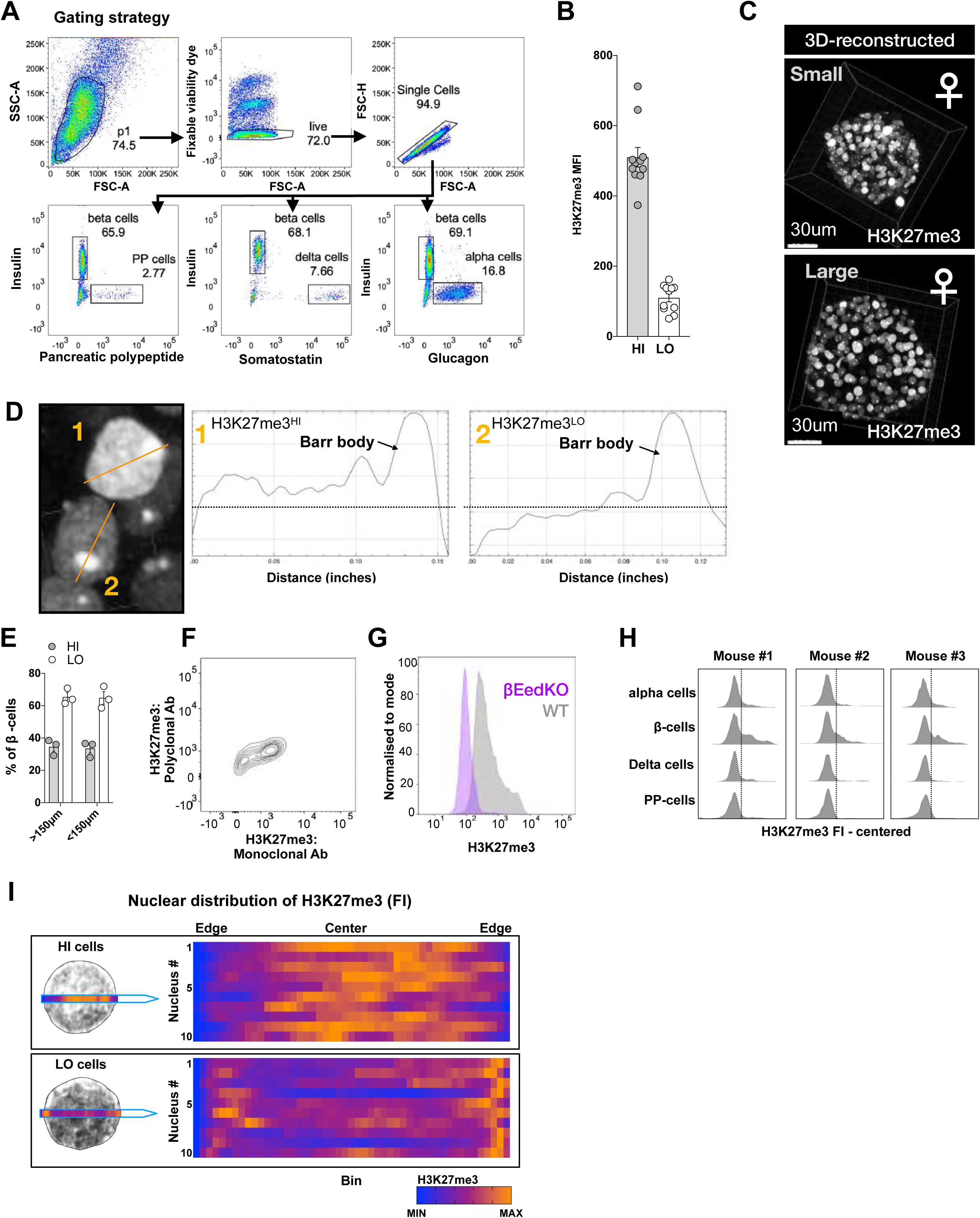
Two epigenetically distinct pancreatic β-cell sub-types. A. Islet cells flow cytometry gating strategy of the different depicted endocrine cells; dead and dying cells were excluded from analysis by labeling with fixable viability dye. Single cells were gated using the pulse geometry gates. B. H3K27me3 FACS intensities in HI/LO cells isolated from n=10 individual mice from 3 independent experiments. C. Representative z-stack-reconstructed images of small or large sized whole islets isolated from female mice. D. Representative 2D image of H3K27me3 labeling of β-cells from female mice and line plots of the center optical plain of H3K27me3-LO (1) or HI (2) β-cells. H3K27me3-silenced X chromosomes also known as the Barr bodies are unchanged. E. FACS quantification of the percentage of single HI/LO β-cells in small (smaller than 150µm) or large (larger than 150µm) islets. n=3 islet isolations, islets were hand-picked according to size under the microscope. F. Co-labeling of H3K27me3 with 2 validated antibodies, either monoclonal or polyclonal. G. Representative histogram of H3K27me3 labeling of single β-cells isolated from WT or β-cell specific EED KO mice. Representative of 3 experiments H. Reproducible H3K27me3 labeling of ɑ, β, δ, and PP-cells, n=3 mice, insulin positive β-cells and the other single hormone positive cells were stained in the same test tube. I. Heatmap representation of the H3K27me3 intensities across the center optical plane (binned) of the HI/LO sorted β-cells averaged in Figure 1H. Each row represents the min max intensities in an individual nucleus (DAPI positive). showing n=10 nuclei.

**Figure S2.**
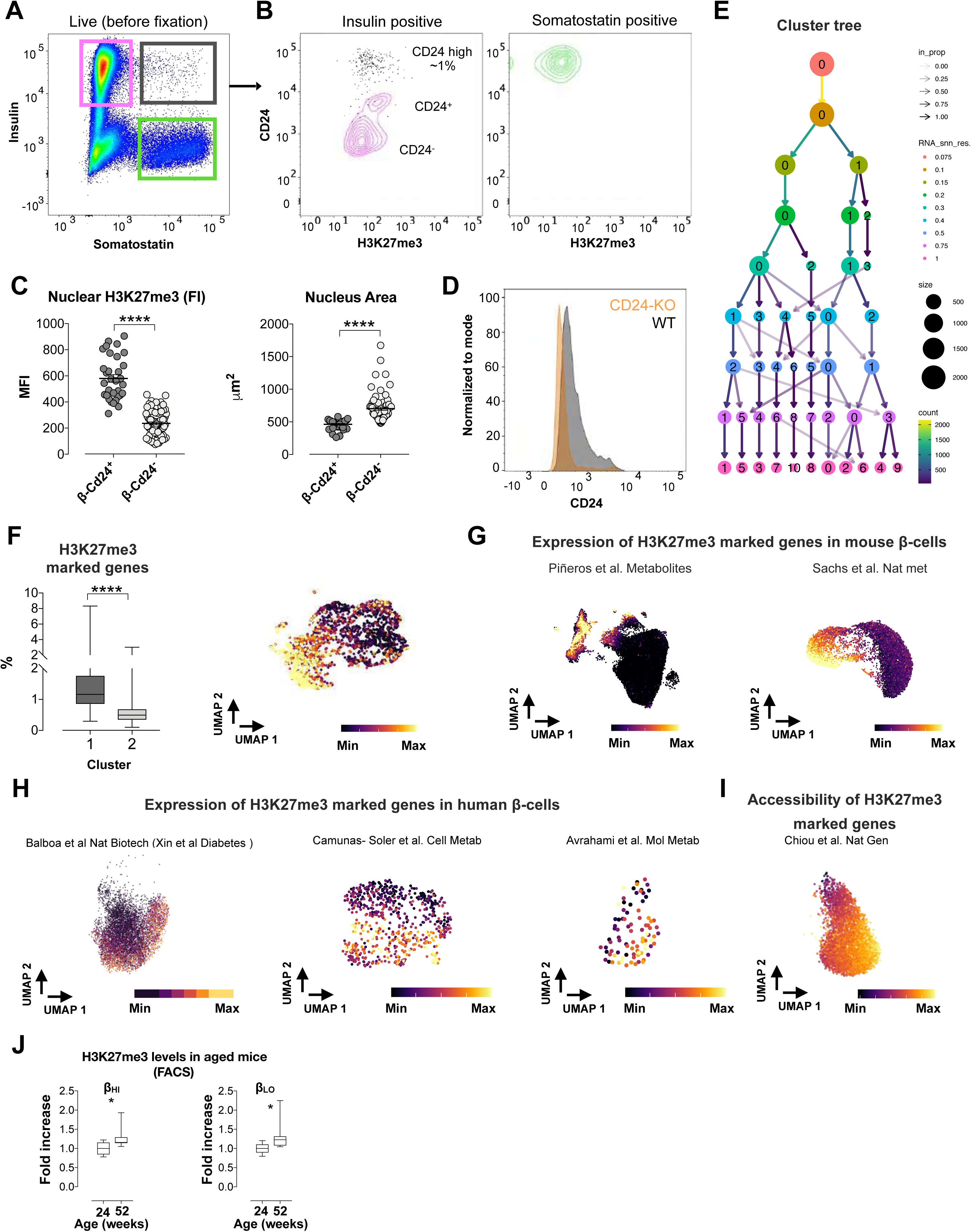
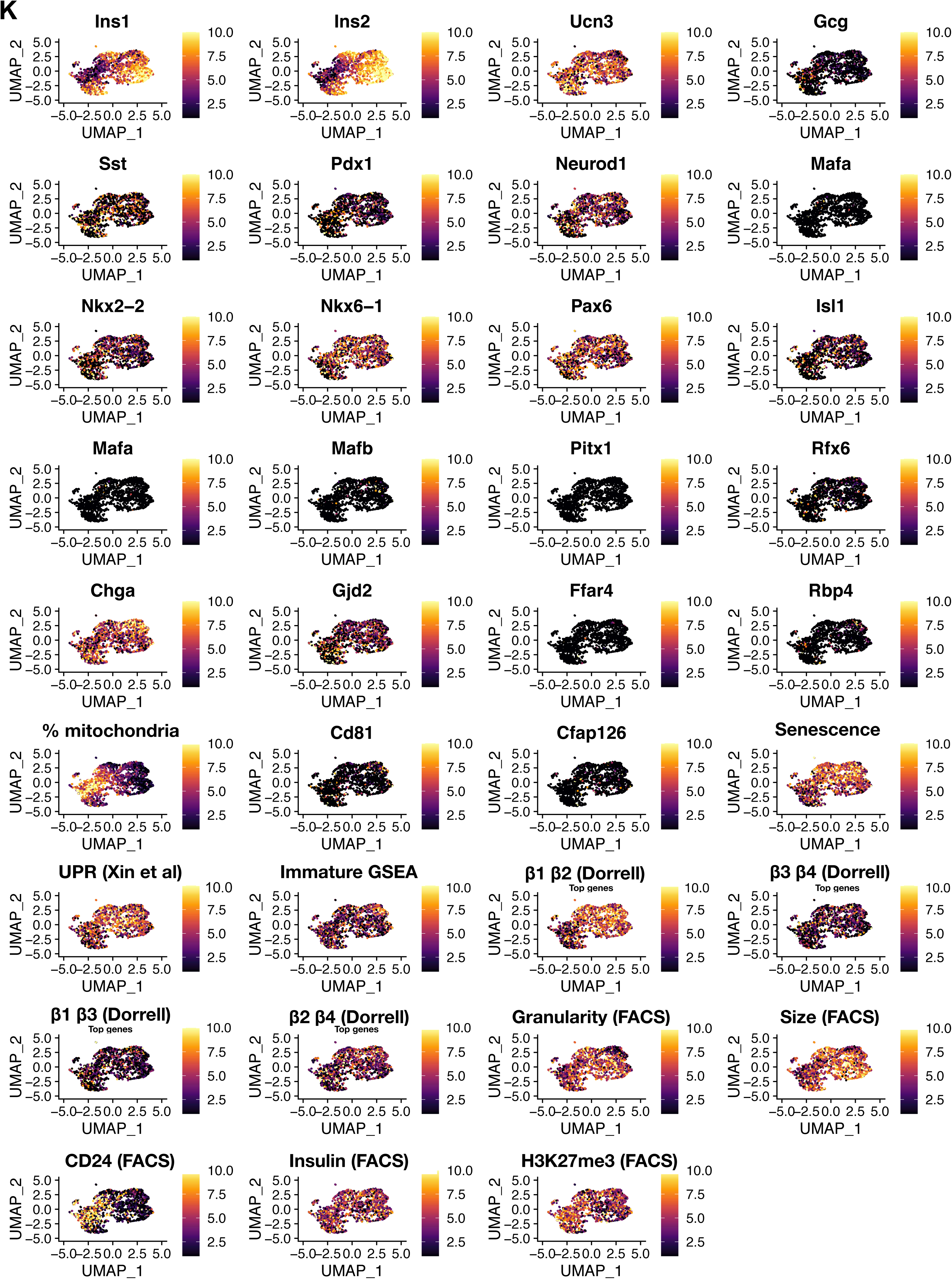
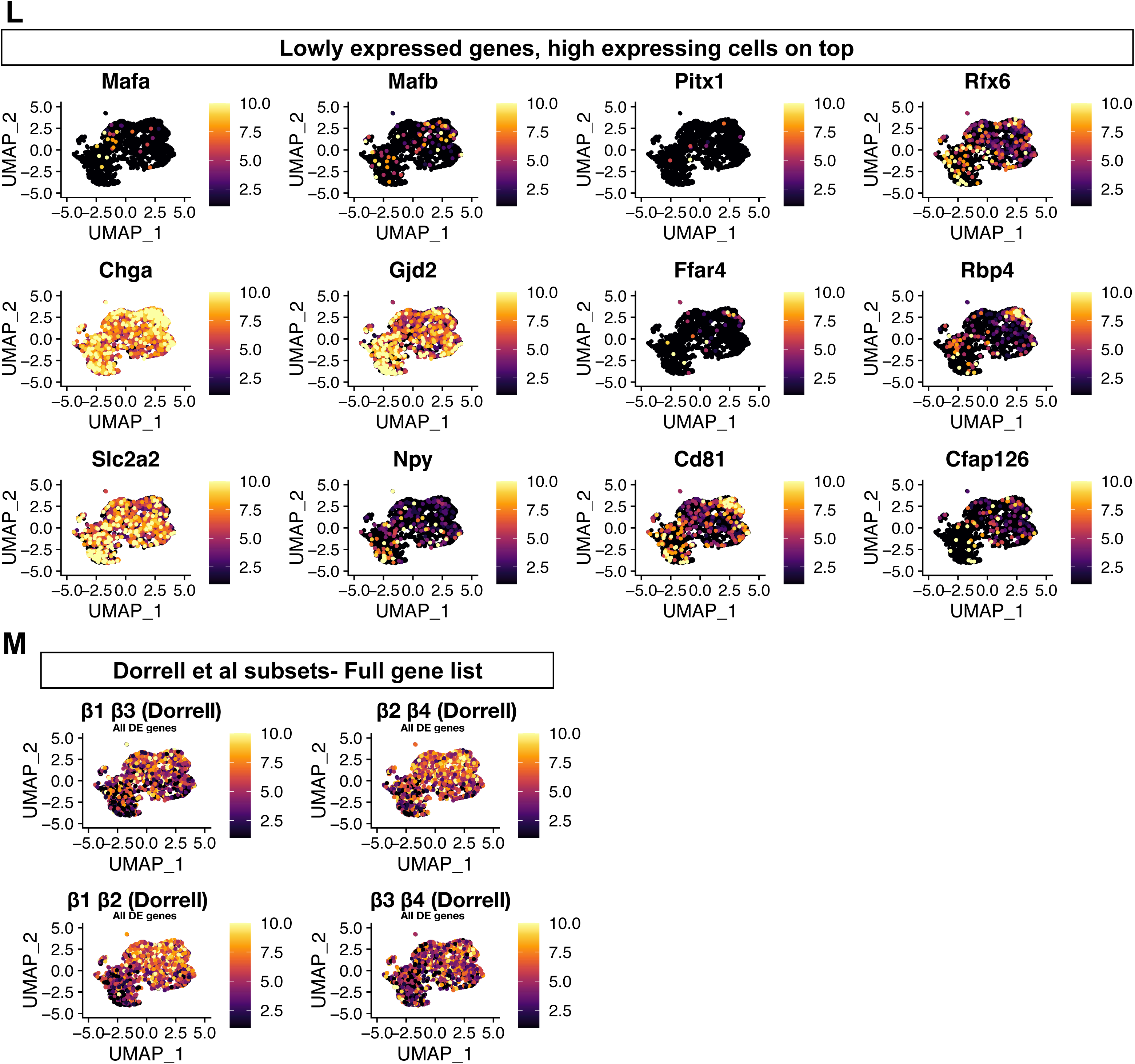
H3K27me3-HI cells are transcriptionally distinct and express cell surface CD24. A. Representative labeling of Insulin and somatostatin in islet cells; gated on cells negative for fixable viability labeling before fixation. Insulin positive β-cells are surrounded pink, somatostatin positive, δ-cells are surrounded green, rare double positive cells are in dark gray. B. Representative H3K27me3 and CD24 labeling in the cells gated above. Insulin / somatostatin double positive cells (dark gray gate) and Insulin positive cells (pink gate) are shown in the left panel; somatostatin single positive cells (green gate) are shown in the right panel. C. Nuclear H3K27me3 levels (MFI) and Nucleus size (area) of CD24^-/+^ β-cells. Cells from β-cell reporter mice were sorted live according to their CD24 levels, fixed and subjected to subsequent staining and confocal imaging. each dot represents a single cell, data are pooled from n=3 individual mice. ****= unpaired t-tests, *p*-value<0.0001. D. Representative labeling of CD24 in insulin positive β-cells isolated from CD24 KO mice compared with their WT littermates. Representative of n=3 experiments. E. Cluster tree visualization of the evaluated Seurat clusters that are determined by the Seurat pipeline at multiple resolutions (RNA_snn_res.). Arrow opacity increase show that low proportion edges appear at higher resolutions, indicating cluster instability. Cluster numbers are determined according to their size and 0 is the largest. Arrow colors and dot size represent the number of cells per cluster. F. Box plot representation of the expression of H3K27me3-marked genes in *SCAN*-seq. The proportions of the expression of H3K27me3 marked genes from all genes per cell per cluster is also overlaid on the UMAP map. ****= unpaired t-test, *p*-value<0.0001. box plots show the median and whiskers indicate min and max values. G. UMAP maps of mouse β-cells from the indicated published data sets overlaid with expression levels of H3K27me3-marked genes each cell. Color coded min to max per experiment. H. UMAP maps of human β-cells from the indicated published data sets overlaid with expression levels of H3K27me3 marked genes each cell. Color coded min to max per experiment. I. UMAP maps of human β-cells from the indicated published data sets overlaid with TSS accessibility levels of H3K27me3 marked genes each cell. Color coded min to max. J. Box plot representation of the fold increase of H3K27me3 FACS levels in aged mice. normalized H3K27me3 levels measured in β_HI_ or β_LO_ cells that were isolated from 24 weeks old or 52 weeks old, aged mice. Isolated from 8-9 mice from n=4 independent experiments. *= unpaired t-test, *p*-value<0.05. box plots show the median and whiskers indicate min and max values. K. UMAP maps highlighting the expression of the indicated gene or sum of expressed genes (Seurat *scaled.data*), Color coded min to max per gene\gene set. L. UMAP maps highlighting the expression of key, lowly expressed gene as in (K), but highest expressing cells are on top. Color coded min to max per gene M. UMAP maps highlighting the expression of all differentially expressed genes as reported in Dorrell et al, 2016.

**Figure S3.**
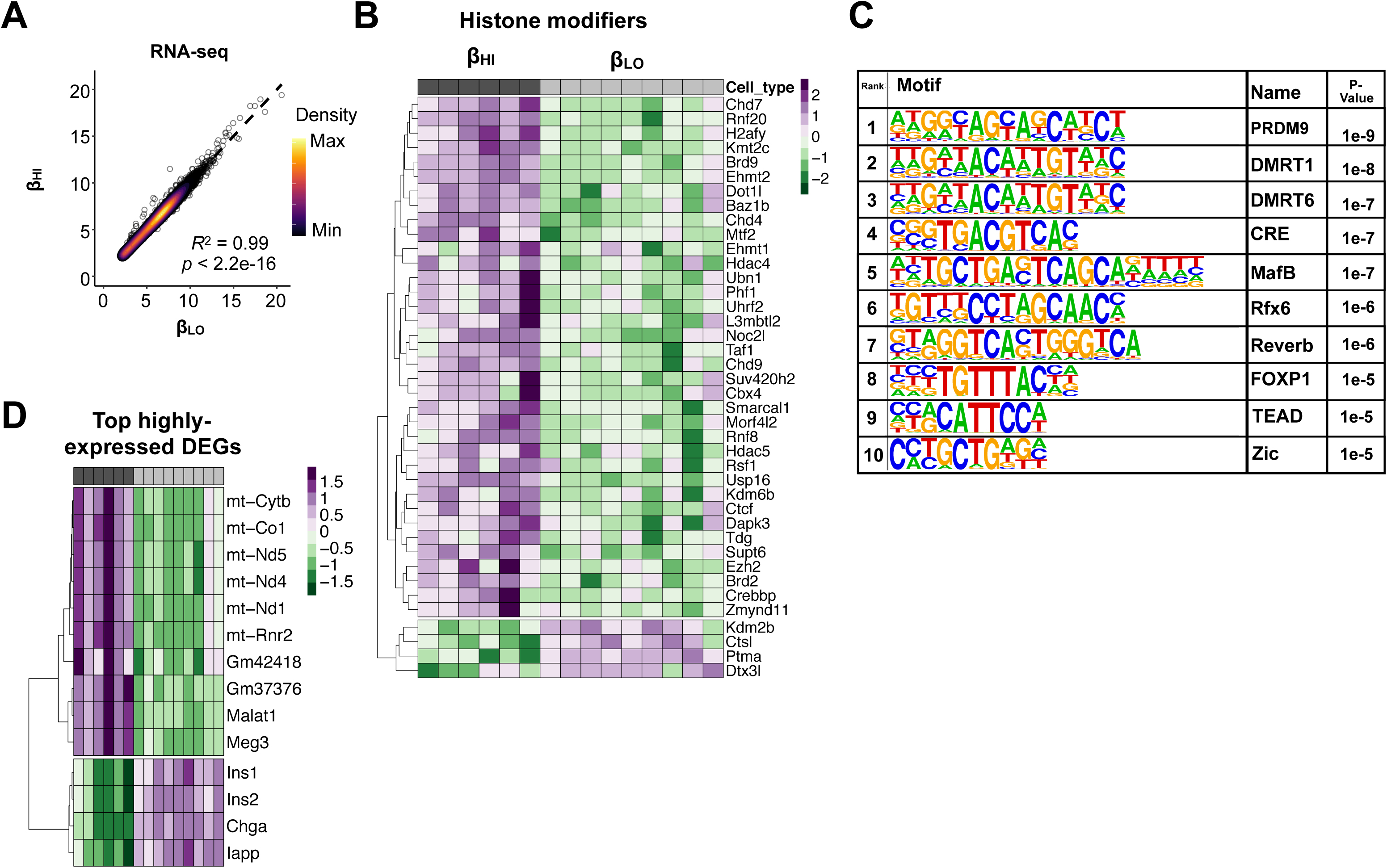
β_HI_ and β_LO_ cells are functionally distinct and specialized. A. Scatterplot showing the correlation of β_HI_ and β_LO_ β-cells’ gene expression profiles. B. Clustered heatmap representation of differentially expressed histone modifiers. Z-score was calculated per gene C. Top 10 significant transcription factor motifs enriched in sequences surrounding the upregulated genes in β_HI_ cells and their P-value. D. Clustered heatmap representation of highly and differentially expressed genes. Z-score was calculated per gene

**Figure S4.**
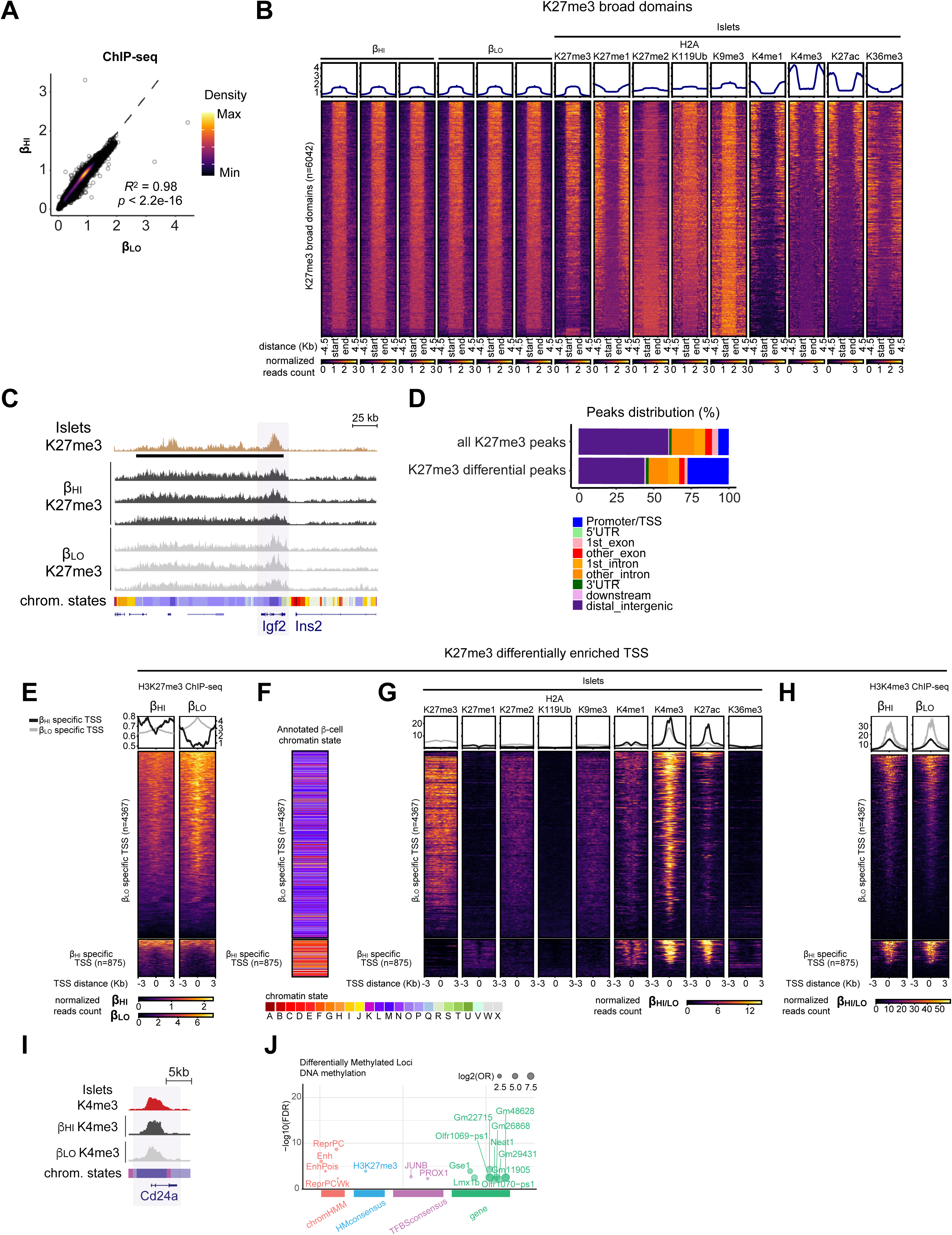
β_HI_ and β_LO_ cells exhibit distinct epigenomes. A. Scatterplot showing the correlation of β_HI_ and β_LO_ β-cells’ H3K27me3 ChIP-seq profiles. Three biological replicates were analyzed for each cell type. B. Heatmap visualization of the indicated histone marks from whole islets, and the H3K27me3 signals from triplicate experiments of β_HI_ and β_LO_ β-cells. The histone marks are visualized on identified K27me broad domains (+/− 4.5 Kb), and show reproducibility among β_HI_ and β_LO_ β-cells replicates, as well as with the K27me3 signal from whole islets. C. Genomic snapshots showing H3K27me3 ChIP-seq tracks from whole islets and β_HI_ and 2 β-cells, as indicated. The INS-IGF2 loci are represented. Horizontal black bars represent H3K27me3 covered broad regions. Colored horizontal bars represent chromatin states, as previously described (Lu et al., 2018), and as indicated in Figure 4G. D. Genomic regions’ distributions among indicated sets of peaks. H3K27me3 differential peaks between β_HI_ and β_LO_ β-cells show a specific enrichment on TSS and a relative depletion of distal intergenic regions, compared to the set of all identified peaks (related to Figure 4E). E. Heatmap visualization of H3K27me3 ChIP-seq signal from merged triplicate experiments of β_HI_ and β_LO_ β-cells, on differentially H3K37me3-enriched TSS between the two β-cells types. H3K37me3 is visualized on TSS +/− 3 Kb. F. Visualization of chromatin states enrichment on differentially H3K37me3-enriched TSS between β_HI_ and β_LO_ β-cells (shown in S4G), show a relative enrichment of bivalent histone decoration (K27me3+K4me3) on β_HI_ β-cells, and of active/transcribed TSS (K4me3+K27ac) on β_LO_ β-cells. Previously annotated chromatin states are reported in figure 4F. G. Heatmap visualization of the indicated histone marks from whole islets, on differentially H3K37me3-enriched TSS between β_HI_ and β_LO_ β-cells. The histone marks are visualized on TSS +/− 3 Kb. H. Heatmap visualization of the H3K4me3 RELACS signals, from merged replicates of β_HI_ and β_LO_ β-cells, on differentially H3K37me3-enriched TSS between the two β-cells types. The H3K4me3 mark is visualized on TSS +/− 3 Kb. It shows no major changes among the two cell types, and a relative enrichment on β_HI_ H3K27me3-enriched TSS. I. Genomic snapshots showing H3K4me3 ChIP-seq tracks from whole islets and purified β_HI_ and β_LO_ cells, as indicated. The Cd24a gene is represented. Horizontal black bars represent H3K4me3 covered broad regions. Colored horizontal bars represent chromatin states, as previously described (Lu et al., 2018) and reproduced in panel (E). J. Enrichment analysis of differentially methylated loci (DMLs) in β_HI_ cells .

**Figure S5.**
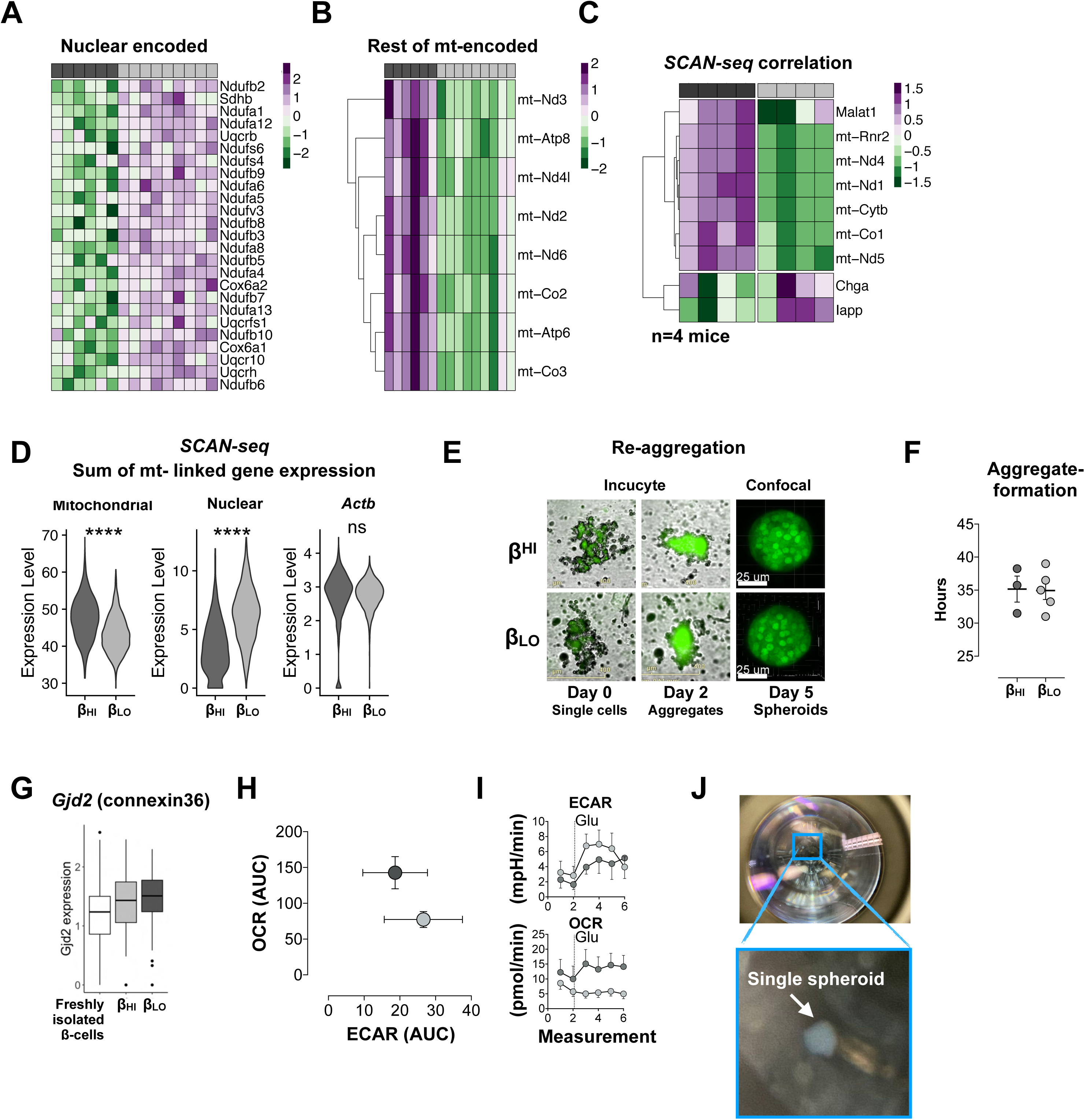
β_HI_ and β_LO_ cells are stably and functionally distinct. A. Clustered heatmap representation of differentially expressed nuclear encoded mitochondrial genes. Z-score was calculated per gene. B. Clustered heatmap representation of differentially expressed mitochondrially encoded mitochondrial genes. Z-score was calculated per gene C. Heatmap representation of *SCAN-seq* scaled and averaged mRNA expression levels of single β-cells positive or negative for CD24 (β_HI_/β_LO_) from 4 individual mice (columns) D. SCAN-seq violin plots of the sum of mt- encoded mitochondrial genes or nuclear encoded mitochondrial genes (unique features) in single β_HI_ or β_LO_ cells. β-actin (*Actb*) expression as house-keeping reference gene. ****= unpaired t-test, *p*-value <0.0001, ns- not significant. E. Representative images of the re-aggregation process of single cells (β_HI_ or β_LO_) isolated form Ins1-YFP reporter mice (green). Single cells (Day 0), visible cell aggregates (Day 2), and spheroids (Day 5). F. The time required for aggregate formation in hours. Each dot represent data from one spheroid. Error bars are mean ± SEM. G. Box plot representation of SCAN-seq derived Gdj2 gene expression in β cells that were freshly isolated or from reaggregated monotypic pseudo-islets. H. Single spheroid metabolic profiling via Seahorse extracellular flux analysis showing total area under the Oxygen consumption rate (OCR) curve and extracellular acidification rate (ECAR) curve presented in Figure S5I. I. Extracellular acidification rate (ECAR) and oxygen consumption rate (OCR) of single pseudo-islets that were given sequential injection of glucose (Glu; 16.7mM end concentration; vertical dotted lines). Presentation of the same data as in Figure 5J. J. Representative image of functionally assayed spheroid (GSIS assay) in a 96 well plate.

**Figure S6.**
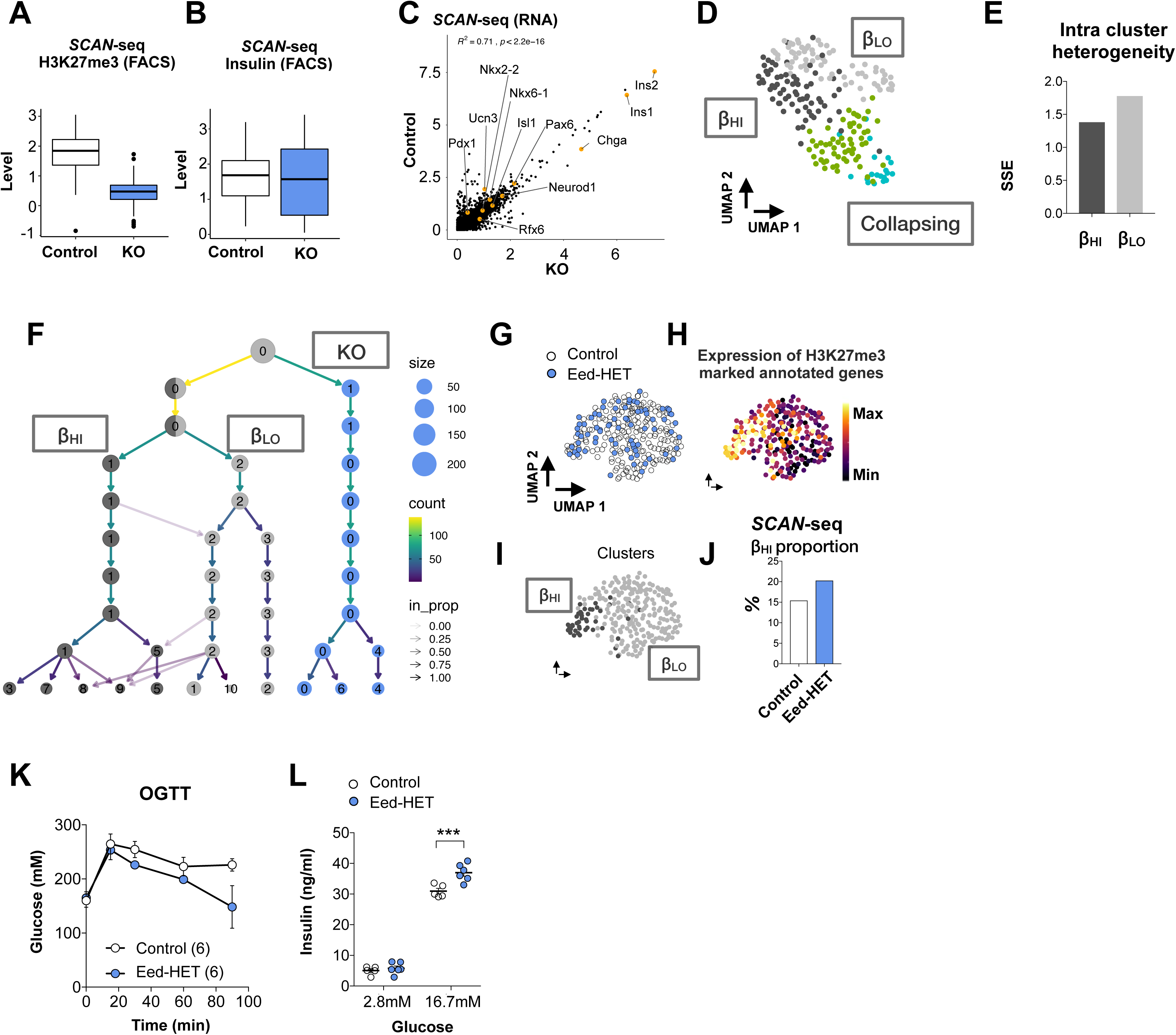
H3K27me3 dosage controls β_HI_ / β_LO_ cell ratios. A. *SCAN*-seq nuclear H3K27me3 levels (FACS intensities) in the β-cells shown in Figure 6A from Control or βEedKO mice. box plots show the median and whiskers indicate the 95^th^ and 5^th^ percentiles B. *SCAN*-seq Insulin protein levels (FACS intensities) in the β-cells shown in Figure 6A from Control or βEedKO mice. box plots show the median and whiskers indicate the 95^th^ and 5^th^ percentiles C. Scatterplot showing the correlation between the SCAN-seq averaged expression profiles (pseudo-bulk RNA-seq) of β_HI_ vs β_LO_ β-cells. Orange labelled dots represent key β-cell genes. D. UMAP visualization of the sub-clustering of the ’A’ β-cell cluster shown Figure 5B. E. Intra-cluster sum of squared errors per the indicated cluster of cells. F. Cluster tree visualization of the evaluated Seurat clusters that are determined by the Seurat pipeline at multiple resolutions. while KO cells build one solid branch (blue), control cells split into two major clusters after which minor clusters emerge. Arrow opacities show that low proportion edges appear at higher resolutions, indicating cluster instability. Cluster numbers are determined according to their size and ‘0’ is the largest. Arrow colors and dot size represent the number of cells per cluster. G. UMAP visualization of sorted mouse β-cells. Colors represent mouse genotypes Eed HET (n=74) or Control (n=178). H. UMAP maps of the β-cells from Figure 5D and 5E overlaid with expression levels of H3K27me3 marked genes. Min to max color coding. I. Cluster topology for the data set in (G) J. Bar plot representation of the percentage of cells that are clustered as β_HI_ cells per genotype in (D) and (E). Colors represent mouse genotypes Eed HET or Control. K. Representative FACS plots displaying H3K27me3 and CD24 labeling profiles in Control (right) or Eed-HET (left) β-cells. The representative histograms along the axes show distributions of H3K27me3 (X) and CD24 (Y). L. Blood glucose levels during oral glucose tolerance test (OGTT; 1 g/kg) in Control or βEed-HET mice, n=6 mice per group. M. Glucose stimulated insulin secretion from whole islets isolated from Control or βEed- HET mice. *** = two-way ANOVA with multiple comparisons correction, *p*-value= 0.001. n=6 replicates, representative of 2 assays.

**Figure S7.**
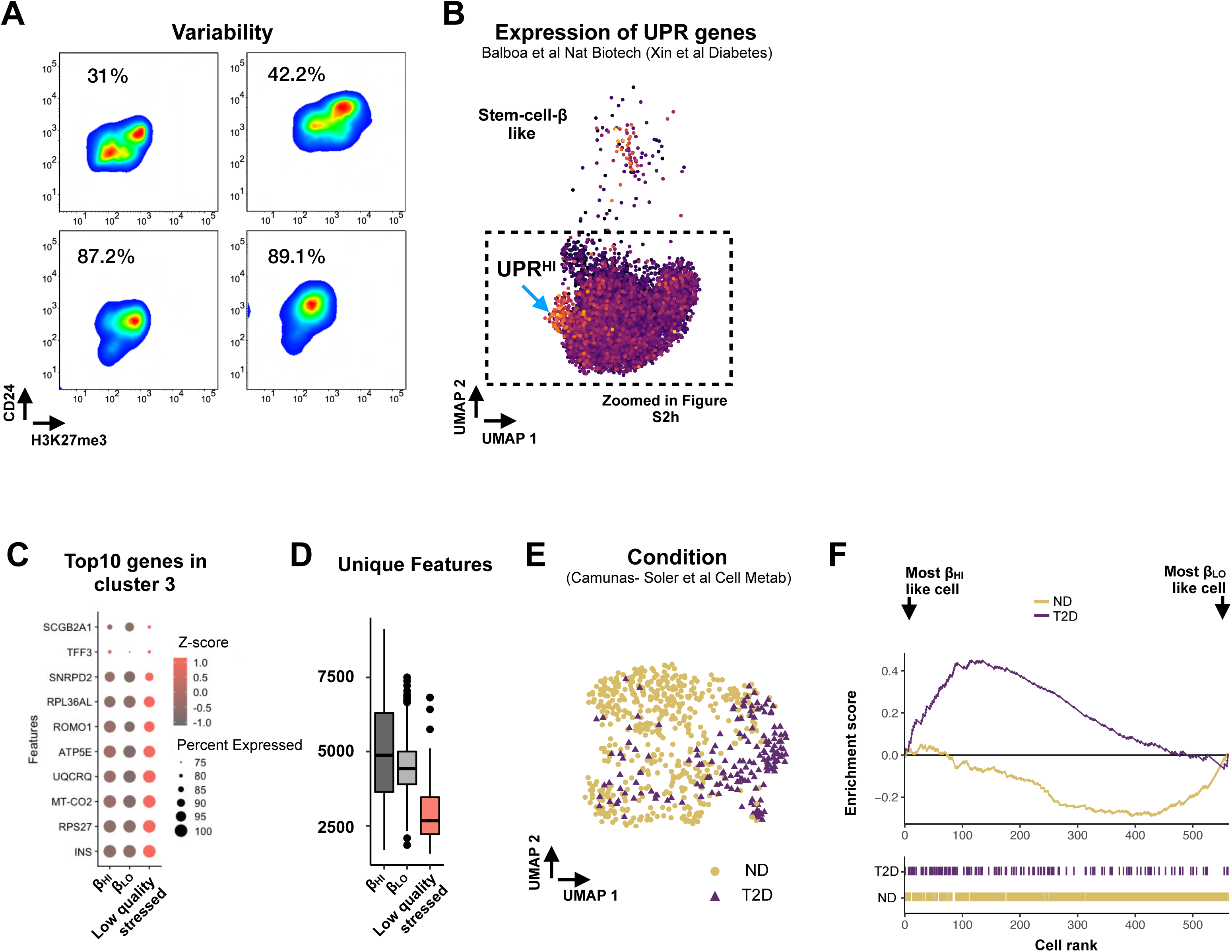
β_HI_ and β_LO_ β-cells are conserved in humans and their ratio altered in diabetes. A. CD24 and H3K27me3 MFIs of insulin positive β-cells (negative for somatostatin, glucagon PP CD31, and CD45) from 4 individual donors. The percentage of CD24 positive, β_HI_ cells is displayed in upper left. For each donor unique gating strategy was applied based on its corresponding unstained sample. B. UMAP maps of all the human β-cells from (Xin et al., 2018) as published in (Balboa et al., 2022). That are also zoomed in Figure S2A, arrow point on UPR high cells, color represent the expression of the UPR related genes *ATF3*, *ATF4*, *DDIT3*, *XBP1*, *CREB3* as reported in (Xin et al., 2018). C. Dot plot representation of the top 10 differentially expressed genes in cluster number 3 across all clusters. Color-code represents average expression (z-scored) and dot size represents the percentage of cells that are expressing the indicated gene per cluster shown in Figure 7D. D. Box plot representation of the uniquely mapped reads (called by Seurat pipeline ‘nFeature_RNA’) per each of the clusters shown in Figure 7D. box plots show the median and whiskers indicate the 95^th^ and 5^th^ percentiles. E. UMAP maps showing the distribution of β-cells from non-diabetic (ND; yellow) and type 2 diabetic (T2D; dark blue). F. Distribution plot of the β_HI_/β_LO_ ranking of single beta cells isolated from ND or T2D donors. Beta cells from T2D are enriched in β_HI_ like signature.

## ChIP-sequencing and RELACS analysis

Mouse H3K27me3 ChIP-seq (n=3 biological replicates for each β-cell type) and H3K4me3 RELACS (n=1 biological replicate for each β-cell type) data were processed and analyzed using snakePipes 2.5 (Bhardwaj et al., 2019) ‘DNA-mapping’ and ‘ChIP-seq’ pipelines. Reads were trimmed and quality controlled using Cutadapt (Kechin et al., 2017) and FastQC (https://www.bioinformatics.babraham.ac.uk/projects/fastqc/), respectively. Mouse reads were mapped with Bowtie2 (Langmead and Salzberg, 2012) on GRCm38/mm10 genome. High quality (MAPQ>3) and properly paired mapped reads were filtered for optical/PCR duplicates using samtools view (Danecek et al., 2021). Coverage tracks for visualization in IGV or UCSC genome browsers were created with the DeepTools (Ramirez et al., 2016) v3.3.2 command ‘bamCoverage’ and normalized to sequencing depth. Spearman correlation matrices of the H3K27me3 signal over the whole genome, among replicates and previously reported ChIP-seq experiments from whole islets (Lu et al., 2018) were generated with the DeepTools commands ‘multiBigwigSummary’ and ‘plotCorrelation’. H3K27me3 peaks and broad domains were called on each single replicate using MACS2 v2.2.6 (Zhang et al., 2008) in ‘broad’ mode and epic2 (Stovner and Saetrom, 2019) v0.0.41 (bin size = 1000, gaps allowed = 10), respectively. PCA on counts over all identified H3K27me3 peaks among all β_HI_ and β_LO_ replicates was performed in R using the command ‘prcomp’. Annotation of identified H3K27me3 peaks according to genomic regions and quantification of tag counts over specific regions were performed using the ‘annotatePeaks.pl’ commands under the HOMER v4.11 suite (Heinz et al., 2010). Differential H3K27me3 enrichment over annotated TSS was performed by running DESeq2 (Love et al., 2014) v1.34.0 on counts tables from biological triplicates. β_HI_ and β_LO_ cell specific TSS were those TSS with H3K27me3 log2(fold change) > 0.2/< -0.2 and adjusted *p*-value < 0.1. Heatmap visualizations and profile plots of H3K27me3 and H3K4me3 signals over specific regions were generated using the DeepTools commands ‘computeMatrix’, ‘plotHeatmap’ and ‘plotProfile’. Chromatin state annotations were based on the previously reported segmentation of the genome from whole islet’s epigenetic landscapes (Lu et al., 2018). Overlaps between annotated chromatin states and genomic regions of interest were found using the ‘intersectBed’ command from bedtools (Quinlan and Hall, 2010). Further analyses (i.e., boxplots, scatterplots) were performed in a R environment using RStudio.

## Quantification of transcript abundance in *SCAN*-seq

Paired end reads were processed using scRNA-seq function in snakePipes (Bhardwaj et al., 2019) (v1.3.0). Briefly, cell barcodes and UMI’s from read 1 were moved into the header of read2 that was then trimmed for adaptors and polyAs using cut adapt (v2.1). The subsequent alignment to the mm10 reference genome (GRCm38) was performed using STAR (v2.4.2a). Raw counts are extracted using feature counts (v1.6.4) using gene annotation version M9 of gencode, and pseudogenes were removed.

## scRNA-seq and SCAN-seq analysis

These data were analyzed using the Seurat v4 algorithm (Stuart et al., 2019), which was followed by standard preprocessing. In brief, cell filtration threshold was set to unique feature counts >700 and >40% mitochondrial genes for *SCAN-*seq or 1000 and >20% mitochondrial genes for the published droplet-based data sets. After QC filtering, the data were normalized by employing a global-scaling normalization method that normalizes the feature expression measurements for each cell by the total expression, multiples the results by 10000 and log-transforms the product. Highly variable transcripts were identified using the Seurat4 FindVaribleFeatures function. The data were scaled while batch correction was applied using the vars.to.regress option to provide an equal weight in downstream analysis and buffer the noise of highly-expressed genes. Then, linear dimensional reduction was applied on the scaled data. Finally, to explore feature expression similarities and defining cell populations, we generated the UMAP using the first 10 principle components/dimensions. Index-sorting files were used to integrate FACS parameters with the seurat object using the CreateAssayObject function. Cluster trees were generated using the ‘clustree’ package. To visualize the expression of groups of genes (Figure S2) a sum of the Seurat *scaled.data* was calculated first. Senesence associated genes (*‘Cellular senescence’*) were called from the mouse Gene Ontology. Genes from the previously reported four β-cell subsets were taken from dorrell et al’s figure 4 (‘top genes’ in Figure S2K) or from the supplemental gene list (All DE geens in Figure S2M). The trajectories in Figure 6B and Figure 7D was generated using slingshot to connect the centroids of each cluster. The code to preprocess and integrate FACS data with the Seurat object will be shared upon request. The custom GSEA in Figure S7 was done based of β_HI_/β_LO_ signature genes (Figure 3C). The mean expression (z-score) for the two gene sets was calculated, then the magnitude and direction of differential signatures was determined by calculating the difference in expression between the two gene sets. The cells were then ranked by difference z-score. All analyses were performed in a R environment using RStudio.

## Analysis of published single-cell/nucleus sequencing

Mouse and human single-cell count matrices from published islet single-cell sequencing datasets were obtained from (Avrahami et al., 2020; Balboa et al., 2022; Camunas-Soler et al., 2020; Pineros et al., 2020; Sachs et al., 2020), scRNA-seq data was preprocessed as described above with the following modifications. cell filtration threshold was set to unique feature counts >1000 and >20%. For the comparisons of cells from ND / T2D donors (Camunas-Soler et al., 2020), data integration (Stuart et al., 2019) was applied using sex as a covariate, only ‘FACS’ cells were analyzed from a total of 11 nondiabetic and 7 T2D donors. Single nucleus ATACseq data (GSE160472) was obtained from (Chiou et al., 2021). Processed and demultiplexed fastq files were reformatted to for subsequent ArchR pipeline, reads were aligned to the reference genome h38 using Chromap with ‘preset atac’ and to produce .sam files that were ultimately converted to Arrow files using the createArrowFiles function with mints = 4 and minFrags = 1000. After quality control replicate 2 and replicate 3 were used. Initial clustering on all islet cells was performed to identify the β-cells that were further used for mapping the sum of expression of H3K27me3 marked genes as previously annotated for the mouse (Lu et al., 2018) or human (Bramswig et al., 2013) data sets.

## Other statistical analysis

Statistical tests for comparisons between conditions in the bar/box plots were performed as indicated in the figure legends, in GraphPad prism v8.

## Data availability

All bulk RNA-seq, scRNA-seq, ChIP-seq/RELACS and DNA methylation array data generated in this study were deposited to Gene Expression Omnibus (GEO) database, and will be publicly available.

## Code availability

The SeSAMe wrapper pipeline SeSAMeStr was published online in zenodo under DOI 10.5281/zenodo.7510575. No other custom code or mathematical algorithm were generated in this report. All publicly available codes and tools used to analyze the data are reported and referenced in the Methods section.

